# FMRP phosphorylation modulates neuronal translation through YTHDF1

**DOI:** 10.1101/2022.11.29.518448

**Authors:** Zhongyu Zou, Jiangbo Wei, Yantao Chen, Yunhee Kang, Hailing Shi, Fan Yang, Shijie Chen, Ying Zhou, Caraline Sepich-Poore, Xiaoxi Zhuang, Xiaoming Zhou, Hualiang Jiang, Zhexing Wen, Peng Jin, Cheng Luo, Chuan He

## Abstract

Translation activation of local synaptic mRNAs is critical to learning and memory^1-3^. Despite extensive studies on how phosphorylation of ribosomal proteins and translation factors enables timely response to exogenous stimuli^4,^ our knowledge on molecular pathways utilized by RNA binding proteins (RBPs) to control translation of specific mRNAs remains incomplete. We have previously found that YTHDF1 regulates depolarization-induced protein synthesis by promoting translation of *N*^6^-methyladenosine (m^6^A)-modified transcripts^5^. Here we report an unexpected mechanism that the stimuli-induced neuronal translation is mediated by phosphorylation of a YTHDF1-binding protein FMRP. Phosphorylation of FMRP serine 499 induced by neuronal depolarization alters the condensing behavior of prion-like protein YTHDF1. Unphosphorylated FMRP sequesters YTHDF1 away from the translation initiation complex, whereas the stimulation-induced FMRP phosphorylation releases YTHDF1 to form translational active condensates with the ribosome to activate translation of YTHDF1 target transcripts. In fragile X syndrome (FXS) models characterized by low FMRP expression, we observed YTHDF1-mediated hyperactive translation, which notably impacts FXS pathophysiology. Developmental defects in an FXS forebrain organoid model could be reversed by a selective small-molecule inhibitor of YTHDF1 which acts by suppressing its condensation in neurons. We characterized transcriptome-wide mRNA translation with inhibitor treatment in organoids and identified targets that explain alleviated FXS pathology. Our study thus reveals FMRP and its phosphorylation as an important regulator of the activity-dependent translation during neuronal development and stimulation, and identifies YTHDF1 as a potential therapeutic target for FXS in which developmental defects caused by FMRP depletion could be reversed through YTHDF1 inhibition.

## FMRP inhibits neuronal translation mediated through YTHDF1

Translation of specific mRNAs is known to be activated upon neuronal stimulation to produce proteins that alter synaptic transmission to facilitate learning and memory^6^. For example, Calcium/calmodulin-dependent protein kinase II (CaMKII) is rapidly synthesized in the dendrites of neurons in the hippocampal CA1 region for long-term potentiation (LTP)^7^. By contrast, elevated Arc/Arg3.1 translation is required for metabotropic glutamate receptor (mGluR)-dependent long-term depression (LTD)^8^. Selective activation of mRNA translation can be achieved through installation of m^6^A marks on specific mRNAs and subsequent recognition by RBPs. YTHDF1 preferentially binds m^6^A-modified transcripts^9, 10^ to promote their translation^11, 12^. In mouse hippocampi, YTHDF1 RNA targets are enriched for transcripts encoding proteins regulating synaptic transmission and LTP. YTHDF1-mediated translation of its target transcripts is not significant at resting state and is activated upon stimulation such as neuronal depolarization with potassium chloride (KCl)^5,13,14^. While this YTHDF1 activation is critical to mouse learning and memory, its underlying regulatory mechanism has yet to be elucidated.

In neurons, YTHDF1 is activated within two hours post KCl depolarization treatment, and then returns to an inactivated state after six hours^5^. This fast and reversible activation suggests a regulation at the post-translational level, which has been implicated in multiple neuronal processes^15^. Therefore, we analyzed global post-translational modification (PTM) changes on cellular proteins upon neuronal depolarization; we specifically examined serine phosphorylation (pS), ubiquitylation (Ub), O-linked β-N-acetylglucosamine (O-GlcNAc), and tyrosine phosphorylation (pY) by western blotting (Extended Data Fig. 1a). Only the global pS level showed a temporal pattern change resembling YTHDF1 activation (Extended Data Fig. 1a), suggesting that serine phosphorylation might be connected with YTHDF1 activation.

As phosphorylation is a key mechanism for protein activation, we examined whether YTHDF1 could be phosphorylated and its phosphorylation may activate its functions in promoting translation. We immunoprecipitated YTHDF1 from mouse brain but failed to detect any noticeable YTHDF1 phosphorylation from these samples (Extended Data Fig. 1b). Therefore, we asked whether any of the potential YTHDF1-binding proteins could be phosphorylated to activate YTHDF1. We isolated protein fractions bound by YTHDF1 upon KCl depolarization in mouse neurons and noticed that a ∼80 kDa protein identified as fragile X mental retardation protein (FMRP) exhibited the most significant increase in serine phosphorylation (Fig. 1a). FMRP is a translation repressor in the brain^16^ known to be associated with m^6^A-methylated transcripts^17^ and YTHDF1^18^. The same interaction between YTHDF1 and FMRP was confirmed in mouse brain tissue (Extended Data Fig. 1c).

**Fig. 1:**
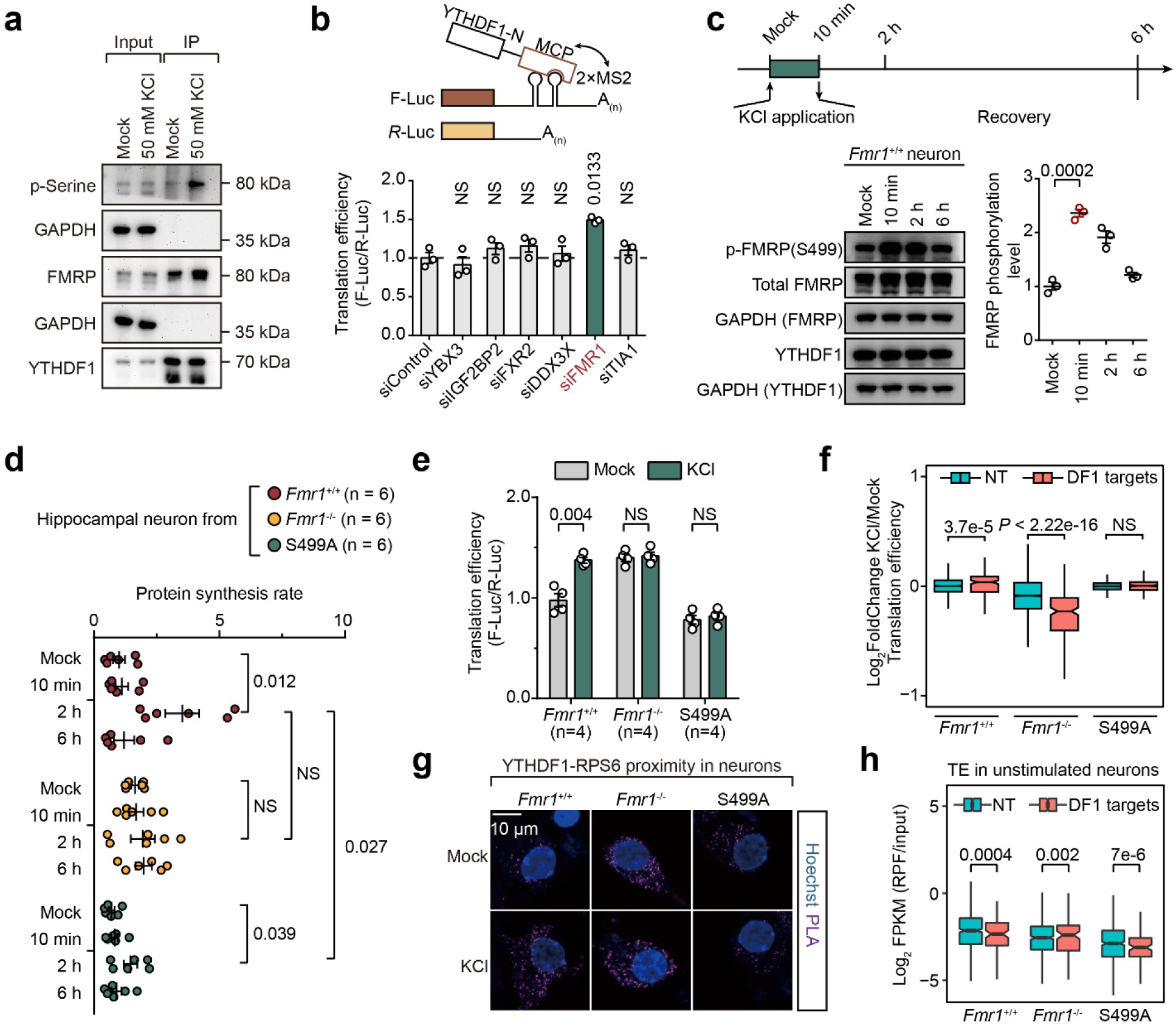
FMRP phosphorylation activates YTHDF1 upon neuronal stimulation. **a**, Western blot analysis of phosphorylated serine in YTHDF1 co-immunoprecipitated fractions. **b**, Reporter assays of YTHDF1-mediated translation in HEK293T cells (n = 3). Proteins negatively regulating m^6^A-mediated translation were knocked-down individually. **c**, Analysis of FMRP phosphorylation upon neuronal depolarization (n = 3 each condition). Western blots are quantified by ImageJ. Relative intensities of different proteins are quantified by normalizing to GAPDH. **d**, Quantification of protein synthesis rate measured by L-Homopropargylglycine (HPG) incorporation in mouse cortical neurons (n = 6 each condition). Fluorescence images are quantified with CellProfiler 3.0 with an in-house pipeline. **e**, Reporter assays of YTHDF1-mediated translation in cultured mouse cortical neurons. **f**, Box plots showing translation efficiency changes of YTHDF1 targets after KCl depolarization in mouse cortical neurons. **g**, Proximity ligation assays in cultured mouse cortical neurons against YTHDF1 and RPS6. **h**, Box plots showing translation efficiency of YTHDF1 targets in neurons at resting state. Data are mean ± s.e.m. Statistical analysis was performed using unpaired two-tailed *t*-test (**b**,**c**,**d**,**e**) or Wilcoxon rank sum test (**f**,**h**). Exact *P* values are indicated, and NS denotes *P* values > 0.05.

In line with its association with YTHDF1, our previous m^6^A-quantitative trait loci (QTL) mapping has also revealed FMRP as one of the top proteins that may suppress translation of the m^6^A-methylated mRNAs (Extended Data Fig. 1d)^19^. To test whether FMRP affects the YTHDF1-mediated translation, we employed a YTHDF1-tethered firefly luciferase (F-Luc) reporter system^11^ and monitored translation changes of this reporter mRNA in HEK293T cells upon individual knockdown of proteins that may suppress translation through m^6^A as revealed from our QTL mapping^19^. To our delight, only when we depleted FMRP translation upregulation of the YTHDF1-tethered reporter was observed (Fig. 1b). Therefore, these observations suggested that FMRP may bind to and sequester YTHDF1 from its translation function.

Serine 499 (S499) in mouse FMRP is known to be a critical phosphorylation site^20, 21^. Therefore, we blotted FMRP phosphorylation with an antibody specific to phosphorylated FMRP S499 at different time points after depolarization. We indeed observed increased FMRP phosphorylation as early as ten minutes post-stimulation in neurons (Fig. 1c). The phosphorylation level of FMRP reduced to basal level after two to six hours, mirroring the temporal pattern of YTHDF1 activation. These results show that FMRP is phosphorylated upon neuronal stimulation and suggest that phosphorylated FMRP may release YTHDF1 for translation promotion.

## Phosphorylation of FMRP regulates translation promotion by YTHDF1

To test whether FMRP phosphorylation mediates YTHDF1 activation during neuronal depolarization, we isolated cortical neurons from *Fmr1^+/+^, Fmr1*^-/-^, or FMRP(S499A), a phosphor-S499 null mutant mice (Extended Data Fig. 1e) and assayed overall protein synthesis rate following KCl depolarization. The basal protein synthesis rate of *Fmr1*^-/-^ neurons is higher than that of *Fmr1*^+/+^ and *Fmr1* (S499A) neurons, consistent with the reported role of FMRP as a translation repressor^16,22,23^. Importantly, while KCl depolarization induced a burst of protein synthesis in *Fmr1*^+/+^ wild-type neurons, neither *Fmr1*^-/-^ nor *Fmr1* (S499A) neurons responded to depolarization (Fig. 1d and Extended Data Fig. 2a), indicating that FMRP S499 phosphorylation is key to the stimulation-induced protein synthesis. Consistently, translation of the YTHDF1-tethered firefly luciferase reporter was elevated in *Fmr1*^-/-^ neurons compared with the wild-type neurons, and depolarization only induced translation upregulation of this reporter in the wild-type neurons (Fig. 1e).

After examining global protein synthesis, we studied the translation efficiency of YTHDF1 targets during neuronal depolarization transcriptome-wide. Translation of YTHDF1 targets was activated after KCl depolarization only in *Fmr1*^+/+^ neurons (Fig. 1f and Extended Data Fig. 3a). *Fmr1*^+/+^ neurons also exhibit the most hyper-translated genes post-depolarization (Extended Data Fig. 3b,c). Correspondingly, YTHDF1 is spatially closer to 40S ribosome component protein RPS6 in *Fmr1*^+/+^ neurons post-depolarization when compared to same neurons without stimulation (Fig. 1g). However, the YTHDF1-RPS6 proximity did not change after depolarization in FMRP(S499A) mutant or *Fmr1*^-/-^ neurons; YTHDF1-RPS6 proximity is always high in *Fmr1*^-/-^ neurons while is always low in FMRP(S499A) mutant neurons (Fig. 1g). Moreover, the translation efficiencies of YTHDF1 targets are higher than non-targets only in *Fmr1*^-/-^ neurons (Fig. 1h), indicating that YTHDF1 is activated in the absence of FMRP. Our results therefore indicate an inhibitory role of unphosphorylated FMRP on YTHDF1 in mouse neurons; FMRP phosphorylation during neuronal depolarization abolishes this inhibition.

## FMRP phosphorylation modulates YTHDF1 condensation

One of the key functions of FMRP S499 phosphorylation is regulating its condensation^24, 25^. YTHDF1 and FMRP are both known to contain intrinsically-disordered domains and prone to form condensates. YTHDF1 is a prion-like protein and proteins containing prion-like domains (PLDs) can fold into distinct structures, which has been implicated in synaptic plasticity formation^26, 27^. To this end, we monitored YTHDF1 condensing behaviors in neurons with immunofluorescence and observed that YTHDF1 foci were formed 10 minutes post-depolarization in *Fmr1*^+/+^ neurons (Fig. 2a). While visible YTHDF1 clusters were observed under both conditions (mock and KCl) in *Fmr1*^-/-^ neurons, the phosphorylation site S499A mutation ablated the foci formation (Fig. 2a, right panel). In an in vitro assay, cellular extracts treated by FMRP kinase inhibitor (Extended Data Fig. 4a, CX-4945 inhibits FMRP phosphorylation at S499^28^) decreased YTHDF1 condensation, as measured by turbidity (Fig. 2b). These data suggest that unphosphorylated FMRP inhibits YTHDF1 condensation.

**Fig. 2:**
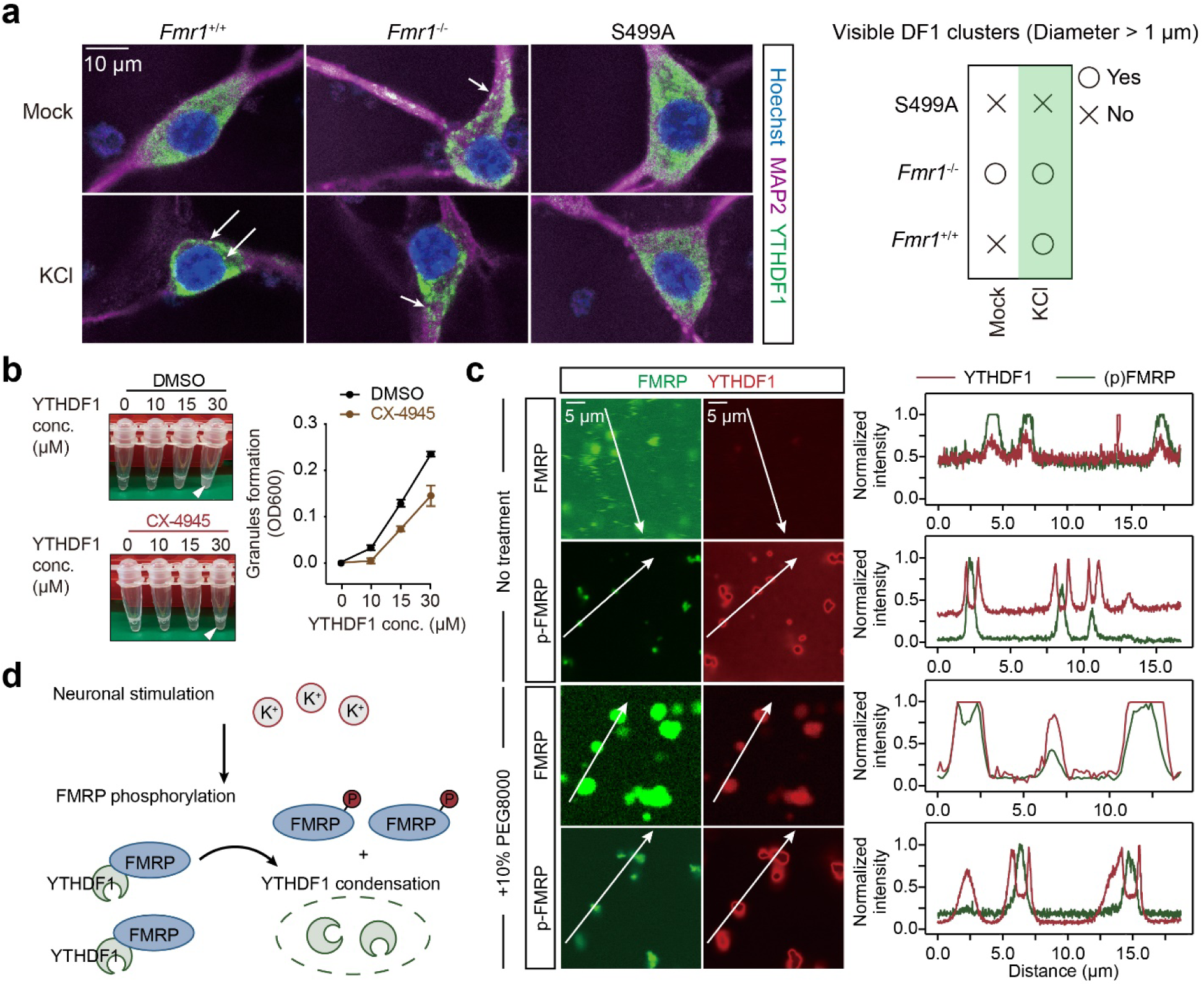
Phosphorylation of FMRP dictates YTHDF1 condensation. **a**, Representative images of DIV 9 cultured mouse cortical neurons labeled by antibodies against YTHDF1 and MAP2. Left, representative images from 3 independent experiments; Right, illustration of condensed YTHDF1 foci formation. **b**, Representative images of purified YTHDF1 incubated with HeLa lysates. YTHDF1 condensation was quantified by turbidity measurement at optical density at 600 nm (OD600). **c**, Representative images of co-condensing of YTHDF1 with unphosphorylated or phosphorylated FMRP (n = 6 each). Scale bar: 40 µm or 5 µm (zoom in). Intensity distribution on a line was quantified by ImageJ: Plot profile module. **d**, Schematic of the activity-dependent FMRP phosphorylation and YTHDF1 condensation.

We constructed an in vitro condensate formation system using only recombinant FMRP and YTHDF1 to study their direct interaction. The addition of unphosphorylated FMRP inhibited YTHDF1 condensation (Fig. 2c). Conversely, YTHDF1 molecules coalesced to form visible condensates under the microscope with phosphorylated FMRP (Fig. 2c and Extended Data Fig. 4b). Moreover, when we induced YTHDF1 condensation by adding crowding reagent PEG8000^29,^ unphosphorylated FMRP colocalizes with YTHDF1 while p-FMRP is mutual exclusive with YTHDF1 (Fig. 2c, lower panel). This suggests a transition in its interaction with YTHDF1 when FMRP is phosphorylated. Additionally, a stronger association between YTHDF1 and FMRP (S499A) compared to FMRP (WT) is revealed by co-IP assay with purified proteins (Extended Data Fig. 4c). This tighter association between YTHDF1 and unphosphorylated FMRP is further supported by increased proximity ligation assay (PLA) signals in cells when treated by CX-4945 (Extended Data Fig. 4d, left panel). Combining all these data, we conclude that unphosphorylated FMRP binds YTHDF1 more strongly than phosphorylated FMRP and this binding represses YTHDF1 condensation, likely with the ribosome translation initiation complex. Indeed, we noticed a significantly decreased spatial proximity between YTHDF1 and a small ribosome unit RPS6 upon inhibition of FMRP phosphorylation with CX-4945 (Extended Data Fig. 4d right panel). Conversely, more YTHDF1 and RPS6 were detected in the RNP granule phase (RG) upon FMRP knockdown using two different siRNAs (Extended Data Fig. 4e). Therefore, unphosphorylated FMRP prevents YTHDF1 condensation, and phosphorylation of FMRP releases this inhibition (Fig. 2d). YTHDF1 condensation in dynamically regulated in neurons during stimulation, and could be linked to mRNA translation.

## YTHDF1 condenses with the small ribosomal subunit to promote translation

To study the functional outcomes of YTHDF1 condensing, we sought to precipitate YTHDF1-containing condensates to reveal condensate-dependent YTHDF1 interaction with cellular biomolecules. According to one report^30,^ condensation of proteins containing PLDs such as YTHDF1 in cultured cells can be triggered by RNase microinjection. We validated that RNase treatment of HEK293T cell lysates induced condensation formation, observed either under a microscope (Fig. 3a) or with turbidity measurements of lysates (Extended Data Fig. 5a). Both 1,6-hexanediol (1,6-HD) and a high concentration of sodium chloride (NaCl) could partially dissolve these condensates, confirming that these condensates are liquid-liquid phase separated granules (Extended Data Fig. 5a). We isolated the RNase-induced condensates from HEK293T lysates using differential centrifugation (Extended Data Fig. 5b) and confirmed that YTHDF1 responds to RNase treatment and is enriched in the RNP granules (RG) (Extended Data Fig. 5c). Of note, 40S ribosome component RPS6 is also significantly enriched in the RG phase after RNase treatment (Extended Data Fig. 5c, left panel). These were also consistently observed in HeLa cells (Extended Data Fig. 5c, right panel). Our data supports that RNase treatment triggers precipitation of RNP granules containing YTHDF1, and thus could be utilized to study its protein interactions.

**Fig. 3:**
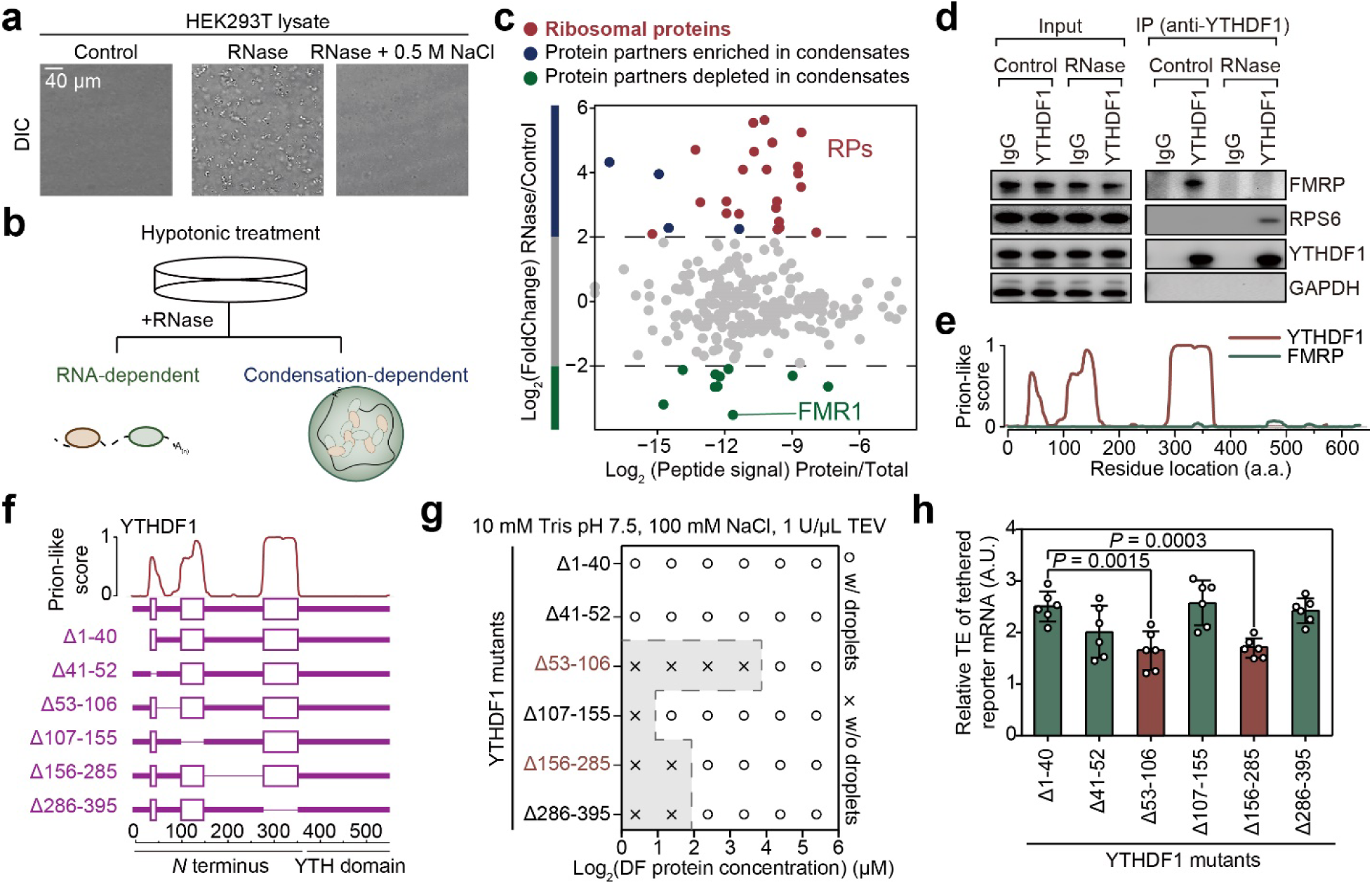
YTHDF1 condenses with the ribosome to promote mRNA translation. **a**, Representative images of HEK293T lysates treated by RNase. Scale bar: 40 µm. **b**, Schematic of using RNase to precipitate prion-like proteins. **c**, Protein partners of YTHDF1 revealed by co-immunoprecipitation with RNase treatment. TMT-enabled multiplexed quantitative MS-Spec data was normalized between two biological replicates. **d**, Western blot analysis of YTHDF1 protein partners with RNase treatment. **e**, Illustration of the prion-like features of YTHDF1 and FMRP calculated by PLAAC. **f**-**g**, Phase separation diagram of YTHDF1 mutants. **f**, Illustration of YTHDF1 mutants with deletion of different domains. **g**, Granule formation propensity of YTHDF1 mutants. **h**, Reporter assay showing effects of YTHDF1 mutants on mRNA translation (n = 6 for each condition). Data are mean ± s.e.m. Statistical analysis was performed using unpaired two-tailed *t*-test. Exact *P* values are indicated, and NS denotes *P* values > 0.05.

We next designed a workflow to study condensation-dependent protein-protein interactions. Cells are lysed with hypotonic treatment to preserve RNP granules and subjected to RNase treatment to precipitate these granules (Fig. 3b). Then immunoprecipitation is performed to study biomolecule interactions of a specific protein. Co-purified proteins are subjected to subsequent analysis including liquid chromatography-tandem mass spectrometry (LC-MS/MS) identification. We applied this approach to study protein-protein interactions involving protein condensates formed by YTHDF1. We identified proteins co-purified by YTHDF1 under RNase treatment, as protein partners involved in YTHDF1 condensate formation should be enriched in the portion precipitated by RNase treatment. We found that FMRP shows significantly decreased binding to YTHDF1 upon RNase treatment (Fig. 3c, highlighted in green), indicating that FMRP is depleted in main YTHDF1 condensates upon RNase treatment. This finding is further validated by western blotting (Fig. 3d) and is consistent with the notion that FMRP protein lacks clear prion-like domains (Fig. 3e).

Notably, we found that condensation-dependent YTHDF1 protein partner repertoire mainly consists of ribosomal proteins (RPs), and is enriched in proteins involved in translation regulation (Fig. 3c and Extended Data Fig. 4d). This may explain the dependency of the YTHDF1-promoted mRNA translation on condensate formation, with YTHDF1 forming an active translation condensate with ribosome to promote the translation of its mRNA targets. To establish causal relationship between enhanced YTHDF1 condensation and mRNA translation efficiency, we constructed a panel of YTHDF1 mutants with deletion of individual domains based on prion-like score predicted by PLAAC (Prion-Like Amino Acid Composition, http://plaac.wi.mit.edu/) (Fig. 3f). We first tested their condensing tendency in vitro. The mutants with individual deletions of amino acid 53-106 (Δ53-106), 156-285 (Δ156-285) or 156-285 (Δ286-395) exhibited lower translation promotion ability compared to wild-type (Fig. 3g). We found that YTHDF1 mutants with impaired condensing ability are less capable of promoting translation of the reporter mRNA it binds (Fig. 3h and Extended Data Fig. 6a). We also utilized an in vitro translation system in rabbit reticulocyte lysates (RRLs) in which we can inhibit YTHDF1 condensation by adding 1,6-hexanediol, an inhibitor of general liquid-liquid phase separation events. YTHDF1 protein condensation was diminished following 1,6-hexanediol treatment in the translation assay (Extended Data Fig. 6b, left panel). The elevated translation efficiency of the YTHDF1-tethered firefly luciferase reporter mRNA was also abolished after 1,6-hexanediol treatment (Extended Data Fig. 6b, right panel). In cell lysates, YTHDF1 distributed in the 40S ribosome fraction decreased upon salt or 1,6-hexanediol treatment (Extended Data Fig. 6c). We therefore conclude that the condensing propensity of YTHDF1 is critical to its translation promotion function.

Collectively, these observations support a working model that the binding of unphosphorylated FMRP to YTHDF1 prevents it from interacting with translation initiation ribosome components, thus inhibiting the YTHDF1-mediated translation. When FMRP is phosphorylated, YTHDF1 is released to form condensates with the ribosome and promotes mRNA translation (Fig. 2d).

## YTHDF1 may induce aberrant translation activation in fragile X syndrome

Fragile X syndrome (FXS) is the most common known cause of inherited intellectual disability and is caused by loss of FMRP protein during brain development^23^. FMRP is primarily thought to be an inhibitor of mRNA translation events. The hippocampal slices from *Fmr1*-KO mice incorporate 15-20% more [^35^S]methionine into nascent peptides than wild-type controls^31^. Consistent with the translational repression role, FMRP was also reported to be associated with stalled polyribosomes^16^. However, studies of FMRP targets also suggested that FMRP may enhance the translation of its target mRNA in certain neurons, because the association of FMRP-target mRNA with ribosome was reduced in FMRP-deficient neurons^32^. Another study conducted in *Drosophila* oocytes indicated that FMRP maintains translation of a subset of long mRNAs^33^. These conflicting roles of FMRP on translation may suggest the involvement of multiple pathways in FMRP-deficient systems; some of these pathways may play more determinant roles in FXS pathophysiology.

We conceived that FMRP deficiency may lead to activation of the YTHDF1-mediated translation in fragile X syndrome (FXS) systems and this pathway could be physiologically relevant in FXS pathophysiology. Aberrantly active translation was implicated in FXS models^16^; restoring normal translation has been explored as a strategy to ameliorate FXS-related phenotypes^34, 35^. Our discoveries of FMRP-regulated mRNA translation activation through YTHDF1 could explain the hyperactive translation in FXS when FMRP is depleted (Fig. 4a). Thus, we decided to develop and test small molecule inhibitors against YTHDF1-m^6^A binding and evaluate their potential for restoring normal translation in fragile X syndrome (Fig. 4a).

**Fig. 4:**
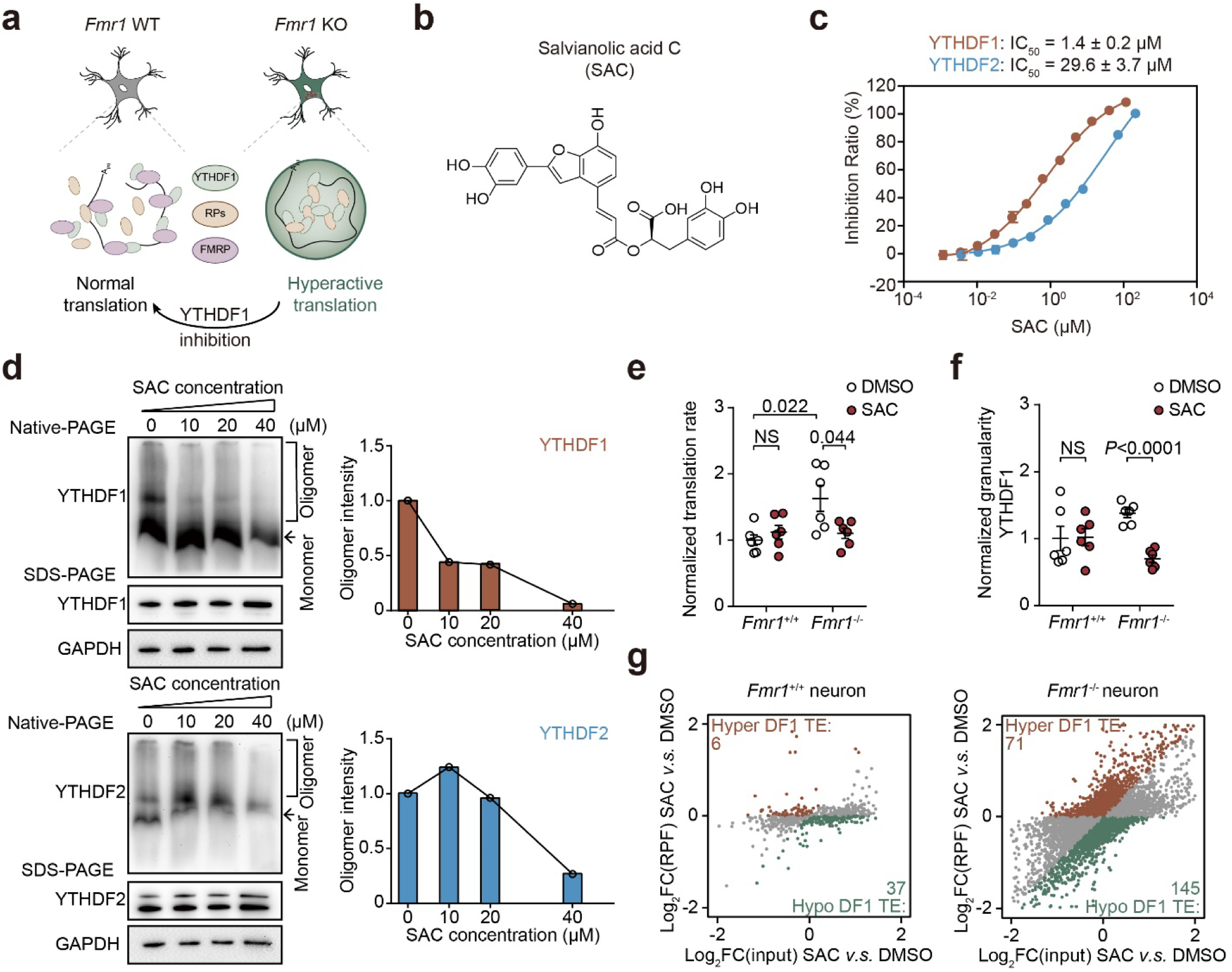
SAC selectively inhibits RNA-binding of YTHDF1 and suppresses YTHDF1 condensation. **a**, Illustration of YTHDF1 inhibition in FMRP-deficient neurons to treat FXS. **b**, Chemical structure of salvianolic acid C (SAC). **c**, IC_50_ value of SAC against YTHDF1 and YTHDF2 measured by AlphaScreen experiments. The selectivity of SAC between YTHDF1 and YTHDF2 is 20:1. **d**, Analysis of YTHDF1 (upper) or YTHDF2 (bottom) protein oligomers in cultured mouse neurons treated with SAC. YTHDF protein oligomers were visualized by Native-PAGE and loading controls were performed with denaturing SDS-PAGE. **e-f**, Quantification of protein synthesis rate (**e**) and YTHDF1 granularity (**f**) in cultured neurons treated with SAC. Protein synthesis and YTHDF1 granularity were quantified with CellProfiler 3.0 with an in-house pipeline. **g**, Scatter plots showing changes in RNA translation in cultured neurons upon SAC treatment. Data are mean ± s.e.m. Statistical analysis was performed using unpaired two-tailed *t*-test. Exact *P* values are indicated, and NS denotes *P* values > 0.05.

## Identification of SAC as a selective inhibitor of YTHDF1

We started with an in-house small molecule library and performed a fluorescence polarization (FP)-based high-throughput screen (HTS) to identify inhibitors of YTHDF1. Salvianolic acid C (SAC) was found to be a potential hit (Fig. 4b). We biochemically confirmed that SAC could effectively disrupt binding of YTHDF1 to its substrate RNA with a half-maximal inhibitory concentration (IC_50_) value of 1.4 ± 0.2 µM in vitro (Extended Data Fig. 7a). We further confirmed the direct binding between SAC and YTHDF1 through nuclear magnetic resonance (NMR) and Carr-Purcell-Meiboom-Gill (CPMG) experiments (Extended Data Fig. 7b). The dissociation constant (K_D_) of SAC against YTHDF1 was determined to be 6.3 µM and 5.3 µM through individual isothermal titration calorimetry (ITC) assay and microscale thermophoresis (MST), respectively (Extended Data Fig. 7c,d). Moreover, AlphaScreen-based substrate competition experiment showed that the IC50 value of SAC against YTHDF1 increased from 1.7 ± 0.2 µM to 46.9 ± 2.3 µM with the addition of unlabeled m^6^A-containing RNA from 0 nM to 400 nM (Extended Data Fig. 7e), suggesting a competitive inhibition but an allosteric inhibition that affects substrate binding is also possible.

Because the YTH domains of YTHDF1 and YTHDF2 share similar sequences, we next investigated the selectivity of SAC against YTHDF1 versus YTHDF2 using the AlphaScreen assay. By comparing IC50 values of the SAC inhibition on individual YTHDF protein interaction with m^6^A-containing RNA (1.4 ± 0.2 µM for YTHDF1 and 29.6 ± 3.7 µM for YTHDF2), we conclude that SAC possesses a decent selectivity against YTHDF1 compared with YTHDF2 (Fig. 4c). PRR5L mRNA, which is known to be a specific target of YTHDF2 but not YTHDF1^36,^ was destabilized only with 50 µM or higher concentrations of SAC (Extended Data Fig. 8b). Translation efficiencies of YTHDF1 target eEF1G and LRPAP1 started to decrease with SAC treatment at 6.25 µM (Extended Data Fig. 8c). Thus, SAC is a specific inhibitor against YTHDF1 and exhibits 20-fold selectivity over YTHDF2. We used 20 µM for most subsequent studies to maximize YTHDF1 inhibition while ensuring YTHDF2 is not affected. This selectivity of SAC against YTHDF1 over YTHDF2 allows us to apply SAC as a tool compound and study biological consequences specifically caused by YTHDF1 inhibition.

## SAC dissolves YTHDF1 condensates and counteracts hyperactive YTHDF1 in neurons

We applied SAC to cultured neuronal cells. Consistent with its ability to block YTHDF1 binding to m^6^A, SAC caused YTHDF1 oligomers disassembling at concentrations starting at 10 µM, while YTHDF2 oligomers were unaffected at SAC concentrations up to 20 µM (Fig. 4d). YTHDF1 and YTHDF2 differ in their N-terminal intrinsically disordered domain sequence (Extended Data Fig. 9a). Indeed, we applied a Thioflavin T (ThT) assay^37^ and found that freshly-purified prion-like domains of YTHDF1 exhibited faster kinetics and higher tendency to form fibrils when compared with that of YTHDF2 within 100-hour monitoring (Extended Data Fig. 9b). In *Fmr1*^-/-^ neurons, SAC treatment restored normal protein synthesis rate (Fig. 4e and Extended Data Fig. 9c) and YTHDF1 granulation (Fig. 4f and Extended Data Fig. 9c) to levels comparable to *Fmr1*^+/+^ neurons. We analyzed translation efficiency of the whole transcriptome and found that SAC treatment caused much more significant alterations in *Fmr1*^-/-^ neurons (Fig. 4g). These data suggest that YTHDF1 is more active in *Fmr1*^-/-^ neurons, with more YTHDF1 targets hypo-translated (145) compared to hyper-translated (71) after SAC-induced inhibition. Together, we demonstrate that SAC not only inhibits RNA-binding of YTHDF1, this inhibition also disrupts YTHDF1 condensation, leading to attenuated translation of target mRNAs of YTHDF1.

## SAC rescues neurodevelopmental deficits in FXS forebrain organoids

We next studied SAC activity on the tissue level in a mouse forebrain organoid system. We have recently established FXS forebrain organoids as a model because behavior phenotypes of FXS can be mild and sometimes challenging to follow. We found that loss of FMRP in human forebrain organoids could lead to reduced proliferation of neural progenitor cells, dysregulated neural differentiation, increased synapse formation, neuronal hyperexcitability, and a deficit in the production of GABAergic neurons^38,^ providing clear phenotypes for us to follow. We developed FXS forebrain organoids from FXS patient-derived iPSCs (induced pluripotent stem cells) and treated both control and FXS forebrain organoids with SAC at 20 µM from day 49 (D49) to 56 (D56) (Fig. 5a). The major immediate outcome of FMRP loss was alterations in mRNA translation, however, it has been challenging to obtain the transcriptome-wide translation profile in human brain. We made use of our human-derived forebrain organoid model and generated the first translation profile, employing a low-input Ribo-seq approach^39^. We identified normal P-site distributions (Extended Data Fig. 10a), validating that our dataset captures translation events.

**Fig. 5:**
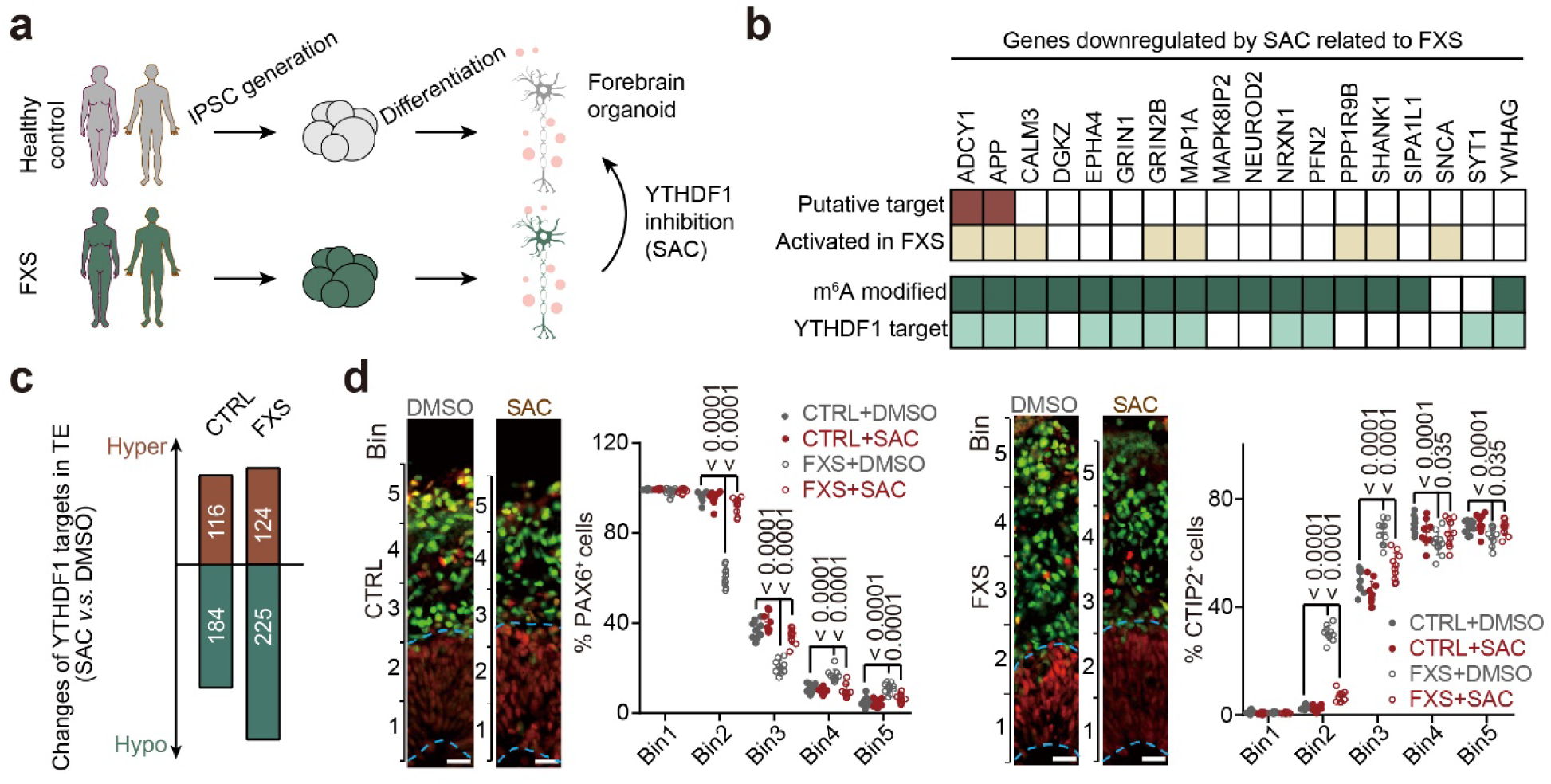
SAC rescues FXS pathology in a human forebrain organoid model. **a**, Illustration of human forebrain organoid model derived from healthy control and FXS patients. **b**, Heat map showing selected downregulated genes by SAC treatment identified in the FXS organoid model. Eighteen genes known to be related to FXS were selected out of 30 downregulated genes enriched in the biological process: regulation of trans-synaptic signaling. **c**, Bar plot showing the numbers of YTHDF1 targets with altered translation in control and FXS organoids upon SAC treatment. **d**, YTHDF1 inhibition rescued deficit of altered neuronal differentiation in FXS forebrain organoids. Shown are sample images (left) and quantification (right) of PAX6^+^ or CTIP2^+^ cells in either control or FXS forebrain organoids under various treatment conditions at day 56. Data are presented as mean ± s.d. (n = 10 sections from at least from 3 organoids each condition. Scale bars: 50 µm. Statistical analysis was performed using unpaired two-tailed *t*-test. Exact *P* values are indicated, and NS denotes *P* values > 0.05.

SAC treatment majorly caused changes in RPFs, instead of mRNA abundance in organoids, showing that YTHDF1 inhibition alters translation efficiency (Extended Data Fig. 10b). We found that the hypo-translated genes, caused by SAC treatment, are enriched in pathways regulating synaptic transmission and plasticity (Extended Data Fig. 10c). Eighteen downregulated transcripts regulating synaptic pathways have been reported to be involved in FXS (Fig. 5b). Most of these transcripts are modified by m^6^A and bound by YTHDF1 (Fig. 5b). Among these genes, *ADCY1* and *APP* are already identified to be therapeutic targets in FXS^40,41,^ supporting our hypothesis that SAC-mediated YTHDF1 inhibition could be applied to alleviate FXS. Similar to observations in neurons, we showed that SAC treatment caused more hypo-translated genes in FMRP depleted FXS organoid (FXS, 225) than control (CTRL, 184) (Fig. 5c).

Next, we assessed the effects of SAC on the neuronal defects in FXS organoids. We found the proliferation deficit in neural progenitor cells (NPCs) was partially rescued by SAC treatment in FXS forebrain organoids by quantifying cells with proliferation marker Ki67 and NPC marker SOX2^42, 43^ detected in immunofluorescence (Extended Data Fig. 11a). A slight increase in the proportion of Ki67^+^ NPCs in control forebrain organoids by SAC treatment was also observed, but not as significant as the effects on FXS organoids (Extended Data Fig. 11a). Moreover, we examined NPC cell cycle kinetics and found that bath application of SAC significantly reverted the phenotype of aberrantly accelerated cell cycle exit in FXS organoids compared to the control (Extended Data Fig. 11b, cell cycle exit was assessed by measuring the proportion of Ki67^-^ and EdU^+^ cells among total EdU^+^ cells within 24 hours’ labeling with EdU labeling). Consistent with the notion YTHDF1 is inactive in control organoids, SAC treatment did not cause a change of cell cycle exit in control forebrain organoids (Extended Data Fig.11b).

To further delineate the effect of SAC on the neuronal differentiation defect caused by the loss of FMRP, we determined the distribution of specific cell types (PAX6^+^ or CTIP2^+^ cells) among equally sized bins spanning the neuroepithelium of forebrain organoids, as previously reported^44^. The number of PAX6^+^ NPCs was significantly reduced in the ventricular zone (VZ)-like layer comprising lower bins while it was increased in the cortical plate layer (higher bins) of FXS forebrain organoids compared to control organoids (Fig. 5d). The alteration in PAX6^+^ NPC differentiation in FXS organoids was rescued after treatment with SAC. CTIP2^+^ cortical plate neurons drastically increased in Bin 2 and 3 in DMSO-treated FXS forebrain organoids compared to DMSO-treated control organoids, illustrating the dysregulated neuronal differentiation and layer specification by FMRP loss (Fig. 5d). When the FXS organoids were treated with SAC, the dysregulated CTIP2^+^ neuronal differentiation and layer organization were rescued to a level comparable to control forebrain organoids (Fig. 5d). Importantly, SAC treatment in control forebrain organoids did not alter CTIP2^+^ neuronal differentiation and layer organization. Consistently, SAC treatment downregulates the translation of key targets such as *ADCY1* and *APP* only in FXS organoids (Fig. 5b).

Together, our results reveal that dysregulated condensation of YTHDF1 and its subsequent translation function upon FMRP deficiency contribute to disease phenotypes in FXS. The small molecule YTHDF1 inhibitor, SAC, could rescue the defects of decreased proliferation of neural progenitor cells, hastened cell cycle exit and altered neural differentiation in FXS organoids by inhibiting the condensing behavior of YTHDF1. Moreover, these rescue effects are specific to FMRP-deficient systems, indicating a dependency on activated YTHDF1. These results further suggest YTHDF1 as a potential target for treating FXS, opening up possibilities for future therapy development (Fig. 5a). Our study also suggests that targeting dysregulated RNP granule equilibrium components may guide drug target discovery in other human diseases such as amyotrophic lateral sclerosis (ALS) and Alzheimer’s disease (AD).

## Discussion

Proteins with prion-like features play important roles in local synaptic mRNA translation, which contributes to synaptic plasticity^27^. Here we report the first example of an activity-dependent phosphorylation of the FMRP/YTHDF1 system as a novel mechanism that neurons utilize to modulate translation in response to stimuli. The phosphorylation of FMRP at serine 499 alters its binding with YTHDF1, a prion-like protein, which causes YTHDF1 condensing with the ribosomal proteins to activate local synaptic mRNA translation. This discovery presents an example of a post-translational modification (PTM) as a switch for segregation of RBPs into different functional RNP granules.

A characteristic feature of fragile X syndrome is FMRP depletion, leading to subsequent aberrant translation. Rescue of aberrant translation has been thought to be a viable strategy to treat FXS^34^. Because FMRP depletion activates the YTHDF1-mediated translation, we reasoned that targeting YTHDF1-mediated translation could re-balance hyper-activated translation and alleviate defects in fragile X syndrome. We developed a selective non-covalent YTHDF1 inhibitor, salvianolic acid C (SAC), and characterized its effect in neurons and in an FXS forebrain organoid model. YTHDF1 inhibition effectively rescued the defects of decreased proliferation of neural progenitor cells, hastened cell cycle exit, and altered neural differentiation in FXS model, further supporting the involvement of YTHDF1 activation in the FMRP-deficient system.

Our study not only suggests YTHDF1 as a potential target in treating FXS, it may also inspire further research on other loss-of-function neuronal diseases or defects related to cytosol RNA processing and translation. Loss-of-function gene perturbations are usually challenging for therapy development because of the difficulty to supply protein agonists. Targeting other RNP granule components to compensate for protein loss could open up various rescue possibilities. With numeral dysfunctions of LLPS events reported in neuronal disorders such as amyotrophic lateral sclerosis (ALS) and Alzheimer’s disease (AD), targeting the dysregulated RNP granules could be a generally applicable strategy towards further understanding of their molecular mechanisms and for developing potential future therapies.

## Main figures

## Methods

### Tethering assay

Tethering assay in HeLa and HEK293T cell lines were performed similarly. About 50 ng reporter plasmid (pmirGlo-Ptight-2BoxB-2MS2) and 250 ng of each effecter plasmid (Flag-MS2, Flag-λ, Flag-YTHDF1N-MS2, Flag-YTHDF1N-λ, or the combination indicated) were used to transfect HeLa or HEK293T in each well of six-well plate at 60%-80% confluency. Six hours later, transfection mixture was replaced with fresh media. Four hours later, each well was trypsin-digested and re-seeded into 96-well plate (1:30) and 12-well plate (1:3) to allow expression of reporter and effector plasmids. Eighteen hours after reseeding, cells in the 96-well plate were analyzed by Dual-Glo Luciferase Assay Systems (E2920, Promega). Firefly luciferase (F-Luc) activity was normalized by renilla luciferase (R-Luc) to evaluate protein production from the reporter. At the same time, samples in the 12-well plate were processed to extract total RNA (DNase I digested) by TRIzol^TM^ reagent (Invitrogen, 15596026), followed by RT-qPCR quantification. The amount of F-Luc mRNA was also normalized by that of R-Luc mRNA. Translation efficiency (TE) of F-Luc mRNA was calculated as the ratio of normalized F-Luc activity (protein level) to normalized F-Luc mRNA level.

Tethering assay in neurons was performed by pre-coating 12-well plates with poly-D-lysine/laminin and seed neurons to coated plates, on day 6 post passaging, cells were transfected with AD1 Primary Cell 4D-nucleofector Y kit (Lonza, V4YP-1A24) with 2 μg reporter plasmid (pmirGlo-Ptight-2BoxB-2MS2) and 16 μg of each effecter plasmid (HA-MS2 or HA-YTHDF1N-MS2) to each well. Fourty-eight hours after nucleofection, neuronal cells are depolarized and collected at different timepoints, luciferase activity was probed with a procedure similar to HeLa and HEK293T cells.

Tethering assay in rabbit reticulocyte lysates was performed by applying in vitro transcribed and capped mRNA encoding mApple-YTHDF1N-MCP or mApple-MCP to the lysates (Promega, L4960) for effector protein translation. After one hour, applying in vitro transcribed and capped mRNA encoding F-Luc-2MS2 and R-Luc were added to the lysates for reporter translation. Luciferase signals were probed after 30 minutes. In vitro transcriptions were performed with a T7 system (Invitrogen, AM1345).

### Polysome profiling

We followed the procedure reported previously^45^. Briefly, cycloheximide (CHX) was added to the media at 100 μ for 7 minutes before lysis and the cell lysate was cleared and digested with Turbo DNase (Invitrogen, AM2238). The cleared lysate was subjected to ultracentrifugation at 28000 rpm for 3 hours at 4 °C and separated on a 5% -50% sucrose gradient.

### Cell fractionation

For RNase treated cell fractionation, one 90% confluent 15-cm dish of WT HEK293T cells was washed with PBS at room temperature once, then harvested with a cell scraper in 2 ml PBS. Cell pellet was resuspended with 3 volumes of lysis buffer (150 mM KCl, 10 mM HEPES pH 7.6, 2 mM EDTA, 0.5% NP-40, 0.5 mM DTT, 1:100 protease inhibitor cocktail, 20 U/ml RNase inhibitor) and incubated on a rotor at 4 °C for 15 minutes. Cell pellet was cleared by centrifuging at 15000 g, 15 minutes, 4 °C. Input samples were saved, and lysates were then separated into two equal volumes, one was subjected to 10 μl RNase A/T1 mix (ThermoFisher Scientific, EN0551), 10 μl lysis buffer was added to the control sample and both samples were incubated at 37 °C for 10 minutes. Then the samples were spined at 4 °C, 15000 g, for 15 minutes. Save supernatant as (Sol) fraction. Resuspend pellet from in 100 μl laemmli buffer (Bio-rad, 1610747) and heat at 90 °C for 5 minutes as (RG) fraction. Load sample for SDS-PAGE followed by western blotting.

### Protein co-immunoprecipitation (co-IP)

Cells, or frozen pellets were lysed in ice-cold RIPA buffer (ThermoFisher Scientific, 89900) with end-to-end rotation for 30 minutes. Then lysates were cleared with centrifugation at 4 °C, 21000 g for 15 minutes. Conjugated protein A&G beads (Invitrogen, 10015D) were applied to the lysate for over night immunoprecipitation. The beads were separated from the supernatant on a magnet during washings with ice-cold RIPA buffer. Final elution was performed by resuspension in 1 x laemmli buffer (Bio-rad, 1610747) and heating at 90 °C for 5 minutes

### RNase-assisted co-IP

For protein co-IP after RNase treatment, one 15-cm dish of confluent WT HEK293T cells was washed once with DPBS. After washing, the cells were harvested in 0.6 ml ice-cold buffer L (50 mM Tris-HCl pH 7.6, 50 mM NaCl, 5 mM MgCl2, 0.1% NP-40, 1 mM DTT, 1x Halt^TM^ protease & phosphatase inhibitor (ThermoFisher Scientific, 78440), 10 U/ml RNase inhibitor (ThermoFisher Scientific, AM2696)) on ice. All subsequent steps were conducted at 25 °C. Cell suspensions were given 30 strokes (1 ml, in 30 s) in a Dounce homogenizer, followed by centrifugation at 2000 g for 2 minutes. The supernatant (Cyt, cytosolic fraction) was collected without disrupting nuclei pellet (Nuc). Cytosol fraction was splitted into two equivalent fractions, 20 μ RNase A/T1 mix (or buffer L) was added to each fraction and incubated at room temperature for 20 minutes while rotation. The two fractions were split to two for IgG and YTHDF1 IP (2 μ each, rabbit IgG isotype control, Abcam ab37415; YTHDF1 antibody, Abcam ab220162) and incubated at room temperature for one hour. Then 10 μl protein A beads (Invitrogen, 1001D) were added to each fraction and tubes were incubated at room temperature for another one hour. Beads were separated on a magnetic device and washed extensively with 1 ml NT2 wash buffer (50 mM HEPES, pH 7.6, 200 mM NaCl, 2 mM EDTA, 0.05% NP-40, 0.5 mM DTT, 200 U/ml SUPERase·In (Invitrogen, AM2694)) at room temperature for 5 times. IP sample was eluted with 80 μ 1x laemmli buffer (Bio-rad, 1610747) at 90 °C for 10 minutes. The eluents were submitted to MS Bioworks for TMT labeling and MS-Spec identification.

### Fluorescence microscopy

For imaging of nascent protein, neuronal cells were depolarized with 50 mM KCl for 10 minutes prior to incubation of L-homopropargylglycine (HPG) for 30 minutes. Then cells were fixed and clicked with Alexa Fluor 488 dye according to manufacturer’s instruction of Click-iT™ HPG Alexa Fluor™ 488 Protein Synthesis Assay Kit (ThermoFisher Scientific, C10428). Samples were imaged on a Leica SP8 laser scanning confocal microscope at University of Chicago. Fluorescence intensity across different samples were quantified with Cellprofiler 3.0 with a custom workflow, total protein synthesis rate was obtained by multiplying average intensity in each cell by the area of each cell. For imaging of protein condensates, the mixtures were prepared in vitro on a 384-well plate with coverslip bottom (Cellvis, P384-1.5H-N). Image acquirement and processing were conducted similarly.

For immunolabeling, neurons were fixed with 4% PFA in DPBS at 37 °C for five minutes, permeabilized with MeOH at -20 °C for eight minutes, dried at room temperature for ten minutes, then washed with DPBS at room temperature for three times. Chambers are blocked in blocking buffer (DPBS, 0.5% BSA, 0.05% Triton X-100, 1:100 SUPERase·In^TM^ (Invitrogen, AM2694)) for one hour at room temperature and YTHDF1 antibody (Abcam, ab220162) was 1:100 diluted in blocking solution and incubate at room temperature for one hour. Chambers were washed with 0.05% Triton X-100 in DPBS for 3 times, then 1:100 diluted goat anti rabbit IgG-AF568 conjugate (ThermoFisher Scientific, A-11011) in blocking solution was added to each well and chambers were incubated at room temperature for one hour. Then chambers were washed with 0.05% Triton X-100 in DPBS for three times and fixed with 4% PFA in DPBS for thirty minutes at room temperature and washed for three times with DPBS. Nuclei was counterstained with 2 µg/ml Hoechst 33342 (Abcam, ab145597) in DPBS at room temperature for 20 minutes, wash with DPBS for 3 times. Chambers were stored at 4 °C before proceeding to imaging.

### Turbidity measurement

Purified YTHDF1 protein was dialyzed to lysis buffer (25 mM Tris-HCl (pH 7.4), 150 mM KCl, 2.5% Glycerol and 0.5 mM DTT). WT HeLa cells treated with DMSO or 2 μ CX-4945 (MedChemExpress, HY-50955) were lysed with lysis buffer on ice and cleared by centrifugation. Different ratios of cell lysates and purified YTHDF1 were mixed and incubated at 37 °C for 10 minutes before analysis of OD600 on a Nanodrop instrument. Turbidity measurements for HEK293T cell lysates were performed similarly.

### RiboLace

Cultured neurons or organoids were treated with 100 µg/ml cycloheximide (MilliporeSigma, C1988) for seven minutes. Cells were spun down and washed once with ice-cold DPBS. Cell pellets were flash-frozen with liquid nitrogen and stored at -80 °C. Total RNA was extracted from cell pellets with standard TRIzol^TM^ protocol. Library constructions were performed with SMARTer Stranded Total RNA-Seq Kit v2 (TaKaRa, 634417) following the manufacturer’s protocols. Ribosome protected fragments (RPFs) were extracted from the cell lysates following the riboLace protocol (IMMAGINA, RL001). RPFs were end-repaired and subjected to library contruction with NEB small RNA kit (NEB, E7560S). Libraries were pooled and sequenced on a NovaSeq6000 sequencer with paired-end mode 50bp setting.

### Quantitative RT-PCR (RT-qPCR)

To quantify expression levels of transcripts, total RNA was reverse transcribed by using PrimeScript™ RT Master Mix (Takara, RR0361) with oligo dT primer and random 6 mers as primer. The cDNA was then subjected to real-time PCR (LightCycler 96 sytem, Roche) by using FastStart Essential DNA Green Master (Roche, 06402712001) with gene specific primers. Relative changes in expression were calculated using the ΔΔCt method.

### Recombinant protein purification

Standard molecular cloning strategies were used to generate N-terminally 6×His tagged prion-like domains of YTHDF1 (residues 69-363), YTHDF2 (residues 69-384) and YTHDF3 (residues 71-390). Recombinant proteins were expressed in E. coli BL21 (DE3) cells grown to OD600 of 0.6 in LB medium. The expression was induced with 0.6 mM IPTG at 16 °C for 20 hours and cells were harvested via centrifugation.

For purification of prion-like domain of YTHDF proteins, cell pellet was resuspended in a lysis buffer containing 25 mM Tris-HCl (pH 7.5), 200 mM NaCl, 20 mM imidazole, 10 mM β-mercaptoethanol (β-ME), and protease inhibitors (Ethylenediaminetetraacetic acid-free protease inhibitor cocktail tablet, MilliporeSigma) and disrupted by sonication for 3 minutes. Cell lysates were clarified via centrifugation at 36000 g for 50 minutes and supernatant was applied to Ni^2+^-NTA resin and washed with lysis buffer, and bound proteins were eluted with lysis buffer supplemented with 300 mM imidazole. The eluted protein was analyzed by SDS-PAGE and concentrated by centrifugal filtration (Amicon Ultra-15). Final concentrated protein was aliquoted, flash frozen, and stored at -80 °C for future use.

### Negative staining transmission electron microscopy

Protein solution (5 μl) of prion-like domain of YTHDF was loaded on an EM grid for 10 seconds and excess solution was removed via blotting with filter paper. The grid was then washed with water and stained with 5 μl uranyl acetate (2%) for 15 seconds. All negative staining samples were imaged on a JOEL 1400 microscopy.

### Fluorescence polarization (FP)

For YTHDF1 inhibitor screening, the compounds were first diluted to 80 μ in the high throughput screening (HTS) experiment. Then the compounds were incubated with 1.25 μ YTHDF1 (aa 361-559) at room temperature for 30 minutes in the assay buffer (20 mM Hepes (pH 7.4), 50 mM NaCl, 0.01% (v/v) tween 20, 5% (v/v) glycerol). And 40 nM 5’-FAM labeled m^6^A-containing mRNA (5’-FAM-UUCUUCUGUGG (m^6^A) CUGUG-3’) was next added and incubated with the mixture at 4 °C for 1 hour before the measurements performed on Envision Readers (PerkinElmer). The fluorescently-labeled m^6^A-containing mRNA was used to adjust the gain factor. The same amount of DMSO and unlabeled m^6^A-containing mRNA were used as the negative and positive control, respectively.

### AlphaScreen

SAC was first diluted from 210 μM to a sequence of concentrations as indicated and incubated with 100 nM His-tagged YTHDF1 (aa 361-559) or His-tagged YTHDF2 (aa 380-579) at room temperature for 30 minutes in the assay buffer (20 mM Hepes (pH 7.4), 150 mM NaCl, 1 mg/ml BSA, 0.01% (v/v) TritonX-100). Then, 10 nM biotinylated m^6^A-containing mRNA (5’-biotin-UUCUUCUGUGG (m^6^A) CUGUG-3’) was added, followed by adding streptavidin donor beads and anti-His acceptor beads in the dark conditions. The samples were kept away from light and incubated at 4 °C for 1 hour. The Alpha signal was detected on Envision Readers (PerkinElmer). The same amount of DMSO was used as the negative control, and the unlabeled m^6^A-containing mRNA (5’-UUCUUCUGUGG (m^6^A) CUGUG-3’) was used as the positive control.

In the substrate competition experiment, unlabeled m^6^A-containing mRNA was first diluted to 400 nM, 300 nM, 200 nM and 100 nM and incubated with 200 nM His-tagged YTHDF1 (aa 361-559) in the assay buffer at 4 °C for 10 minutes. SAC was next diluted from 210 mM and incubated with samples in the assay buffer at room temperature for another 30 minutes. And 20 nM biotinylated m^6^A-containing mRNA was then added to samples. The following steps are as same as aforementioned.

### NMR

SAC was dissolved in 5% deuterated DMSO to the concentration of 200 μ YTHDF1 (aa 361-559) was diluted to 20 μ 10 μ and 5 μ in phosphate buffer (20 mM NaH_2_PO4, 20 mM Na_2_HPO_4_, 150 mM NaCl, pH 7.4, D_2_O). NMR CPMG experiment was conducted at 25 °C using Bruker Avance III spectrometer (600 MHz proton frequency) with a cryogenically cooled probe (Bruker biospin, Germany). Through the pulse sequence (RD−90°−(τ−180°−τ n−ACQ), the solvent-suppressed 1D ^1^H CPMG was obtained. And a 54.78 dB pulse in 4 s duration of the recycle delay (RD) was applied to eliminate water resonance in the presaturation procedure. Then, the 90° pulse length was modulated to 11.82 μ approximately. And at last, a total of 4 dummy scans and 64 free induction decays (FIDs) were collected into 64000 acquisition points, covering a spectral width of 12 kHz (20 ppm) with the acquisition time (ACQ) of 2.73 s.

### Isothermal Titration Calorimetry (ITC)

Freshly purified YTHDF1 (aa 361-559) was first dialyzed at 4 °C overnight and diluted to 50 mM with dialysis buffer (20 mM Hepes (pH 7.4) and 200 mM NaCl) before loaded into the sample cell. SAC was then dissolved and diluted to 1 mM with dialysis buffer and loaded into the syringe. After one 0.4 μ injection, through titrating SAC into YTHDF1 (aa 361-559) by nineteen 2 μ injections at 180 s intervals, the isothermal titration calorimetry (ITC) experiments were performed on the Microcal ITC 200 isothermal titration calorimeter instrument (GE Healthcare) at 25 °C. In order to exclude the thermal effect of background dilution, 1 mM SAA titrating into dialysis buffer was also performed as the control. The experimental data was analyzed using Microcal ORIGIN (v7.0) software (Microcal Software).

### Microscale thermophoresis (MST)

Microscale Thermophoresis (MST) experiments was performed on Monolith NT. Automated instrument (NanoTemper Technologies) via the label-free method. SAC was diluted from 250 μ M purified YTHDF1 protein at the proportion of 1:1. Samples were incubated at room temperature for 20 minutes in MST buffer (20 mM Hepes, pH 7.4, 200 mM NaCl, 0.1mM Pluronic^®^ F-127) and centrifuged at 13000 rpm for 10 minutes at 4 °C before the measurement. Then the samples were loaded into Monolith NT. Automated LabelFree Premium Capillary Chips (NanoTemper Technologies) and the measurements were started. MO. Affinity Analysis Software v2.3 (NanoTemper Technologies) was used to analyze the data and get the K_D_ value of SAC.

### Molecular docking

Molecular docking was performed via Maestro (v17.1. Schrodinger, LLC). The structures of YTHDF1 (PDB ID: 4RCJ) and YTHDF2 (PDB ID: 7Z26) were first prepared using protein preparation wizard module of Maestro, respectively. Then the grids were generated on the basis of the results. Next, the structure of SAC was prepared through ligand preparation module and docked into YTHDF1 and YTHDF2 via glide docking module in the XP mode, respectively.

### Human forebrain-specific organoid cultures

The human iPSC lines were reprogrammed from skin biopsy samples of three FXS male patients and three age-matched healthy males at Emory University. All iPSC lines were cultured on irradiated MEFs in human iPSC medium consisting of D-MEM/F12 (Invitrogen), 20% Knockout Serum Replacement (KSR, Invitrogen), 1X Glutamax (Invitrogen), 1X MEM Non-essential Amino Acids (Invitrogen), 100 µM β-Mercaptoethanol (Invitrogen), and 10 ng/ml human basic FGF (PeproTech). Forebrain-specific organoids were generated using established protocols as previously described^43^. Briefly, human iPSC colonies were detached from the MEF feeder layer with 1 mg/ml collagenase treatment for 1 hour and suspended in embryonic body (EB) medium, consisting of FGF-2-free iPSC medium supplemented with 2 µM Dorsomorphin and 2 µM A-83 in non-treated polystyrene plates for 4 days with a daily medium change. On days 5-6, half of the medium was replaced with induction medium consisting of DMEM/F12, 1X N2 Supplement (Invitrogen), 10 μg/ml Heparin (MilliporeSigma), 1X Penicillin/Streptomycin, 1X Non-essential Amino Acids, 1X Glutamax, 4 ng/ml WNT-3A (R&D Systems), 1 μM CHIR99021 M SB-431542 (Tocris). On day 7, organoids were embedded in Matrigel (BD Biosciences) and continued to grow in induction medium for 6 more days. On day 14, embedded organoids were mechanically dissociated from Matrigel by pipetting up and down onto the plate with a 5 ml pipette tip. Typically, 10-20 organoids were transferred to each well of a 12-well spinning bioreactor (SpinΩ containing differentiation medium, consisting of DMEM/F12, 1X N2 and B27 Supplements (Invitrogen), 1X Penicillin/Streptomycin, 100 µM β-Mercaptoenthanol (Invitrogen), 1X MEM NEAA, 2.5 μ l Insulin (MilliporeSigma). All media was changed every other day. Control or FXS forebrain organoids were treated with a newly developed DC-Y22 (SAC) at 20 μ from day 49 to 56, fixed at day 56, and stained for immunocytochemistry.

### Immunocytochemistry

For immunocytochemistry, forebrain organoids were processed at day 56 as previously described^43^. Briefly, whole organoids were fixed in 4% Paraformaldehyde in Phosphate Buffered Saline (BPS) for 30-60 minutes at room temperature. Organoids were washed 3 times with PBS, incubated in 30% sucrose solution overnight, embedded in tissue freezing medium (General Data) and sectioned with a cryostat (Leica). For immunostaining, freezing medium was washed with PBS before permeabilization with 0.2% Triton-X in PBS for 1 hour. Tissues were then blocked with blocking medium consisting of 10% donkey serum in PBS with 0.1% Tween-20 (PBST) for 30 minutes. The following primary antibodies were used: anti-Ki67 (mouse; 1:500; BD-Pharmingen), anti-SOX2 (goat; 1:500; R&D Systems), anti-PAX6 (rabbit; 1:500; Thermo fisher), anti-CTIP2 (rat; 1:400; Abcam). Primary antibodies diluted in blocking solution were applied to the sections overnight at 4 °C. After washing with PBST, secondary antibodies diluted in blocking solution were applied to the sections for 1 hour at room temperature, washed with PBST and stained with DAPI. For EdU labeling, forebrain organoids were pulsed labeled with EdU (10 µM) for 24 hrs at day 56, followed by fixation and immunostaining. All images were captured by Nikon Eclipse Ti-E microscope. Quantitative analyses were conducted on randomly selected cortical structures in a blind fashion. The numbers of cells positive for each marker were measured using ImageJ software (NIH). To quantify the distribution of specific cell types in the neuroepithelium, the entire span of the neuroepithelium was divided into five equal portions (bins) and measured the distributions of specific cell types by calculating the percentage of each marker in the total DAPI^+^ cells in each bin.

### Primary cortical neuron culture

For the isolation of primary neurons from P1-3 wildtype, Fmr1 knockout or Fmr1 S499A mutant mouse, Papain dissociation system from Worthington Biochemical Corporation (LK003160) was used. Briefly, the whole cerebral hemispheres were dissected out, slightly minced and dissociated in papain dissociation solution containing 20 units/ml papain and 0.005% DNase with constant agitation on a rocker platform for 30 minutes at 37°C. Then, the dissociated tissue mixture was triturated with 10ml pipette. The undissociated tissue was allowed to settle down to the bottom of the tube. Only the cloudy cell suspension was centrifuged at 300g for 5 minutes at room temperature. The pelleted cells were resuspended in reconstituted albumin-ovomucoid inhibitor solution. The discontinuous density gradient was made with albumin-inhibitor solution. The cell suspensions were carefully layered on top, then centrifuged at 70g for 6 minutes at room temperature. The dissociated cells pelleted at the bottom of the tube, and the membrane fragments remained at the interface. The supernatant was discarded and pelleted cells were immediately resuspended in the medium for cell culture or frozen in Bambanker cell freezing media (bulldog bio BBO1) for storage freezing at the rate of -1 °C/minute.

### RiboLace analysis

Adaptors for input and ribosome-protected fragments (RPFs) reads were trimmed with cutadapt (version 1.14)^46^ with the minimum length set to 1 (--minimum-length 1 and --pair-filter any), respectively. The trimmed reads were aligned to the hg38 or mm10 genome^47^ with STAR (version 2.5.2b)^48^ with the following parameters (--quantMode TranscriptomeSAM -- alignEndsType Local). Then the output SAM files were converted to BAM files with samtools (version 1.6)^49^. Posterior_mean_estimate counts (PMC) values for each transcript were calculated with RSEM (version 1.2.28)^50^. Translation efficiency was determined by normalizing PMC of RPFs to input.

## Acknowledgements

This work was supported by NIH HG008935 (C.H.). NS111602, MH116441 and HG008935 to P.J., SAC inhibitor discovery was supported by National Natural Science Foundation of China (21820102008 to H.J. and C.H., 91853205, 81625022, 81821005 to C.L.) and the Science and Technology Commission of Shanghai Municipality (18431907100 to H.J. and 19XD1404700 to C.L.). C.H. is an investigator of the Howard Hughes Medical Institute. We thank Ms. Lindsay Scarpitta and Genomics Facility of the University of Chicago for generous help with high-throughput sequencing. We thank Ms. Linda Zhang for helping with SAC characterization in cell lines and Mr. Ge Sun for molecular docking. We thank the Integrated Light Microscopy Core at the University of Chicago for providing imaging tools and Dr. April Pawluk from LifeScienceEditors for editing.

## Author contributions statement

Conception, C.H. and Z.Z.; experiments, Z.Z., J.W. with help from F.Y.; sequencing data analysis, Z.Z.; organoid culture and experiments, Y.K., Y.Zhou, Z.W. and P.J.; inhibitor development and characterization, Y.C., S.C., H.J. and C.L.; YTHDF1 phosphorylation in mouse brain, H.S.; ThT assay, X.Z.; writing, Z.Z., C. S.-P., Y.K., P.J. and C.H. with input from all authors; supervision, C.H..

## Competing interests

C.H. is a scientific founder and a scientific advisory board member of Accent Therapeutics, Inc., Aferna Green. Inc., and AccuraDX Inc. The other authors declare no competing interests.

## Data availability

High-throughput sequencing data can be accessed in the Gene Expression Omnibus under accession number GSE214882 (Token: sbahyyaephajpux). Source data for all graphs in Figure and Extended Data Figures are provided in separate excel files. Uncropped gel or membrane scans with size marker indications are provided in supplementary figure 1.

---

**Extended Data Fig. 1:**
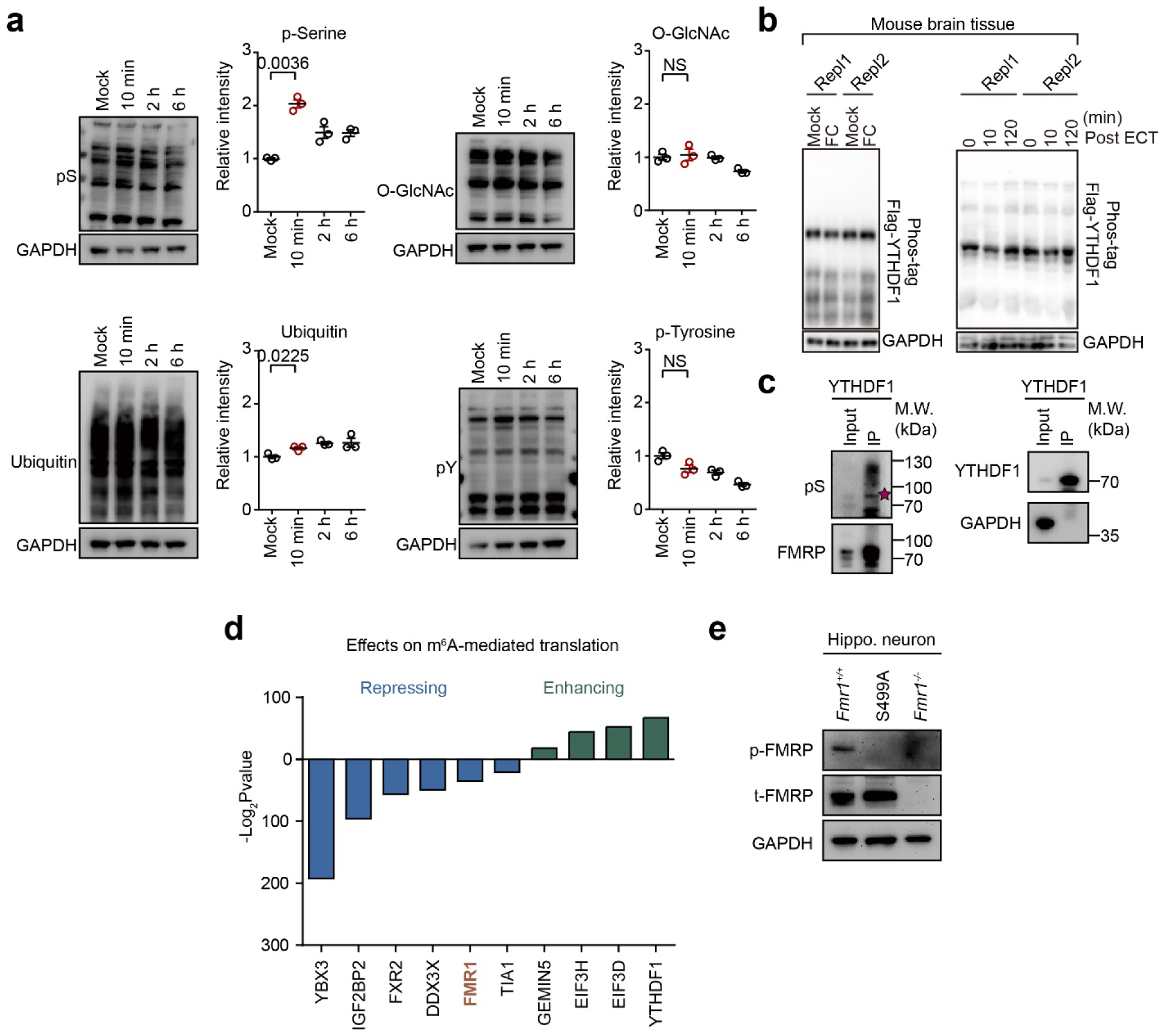
FMRP is phosphorylated at S499 upon neuronal depolarization. **a**, Representative western blots of global post-translational modifications upon neuronal depolarization (n = 3 each condition). Western blots are quantified by ImageJ. Relative intensities of different proteins are quantified by normalizing to GAPDH. **b**, Phos-tag gels analyzing YTHDF1 phosphorylation in the mouse brain after fear conditioning (FC) or electroconvulsive treatment (ECT). **c**, Western blots validating YTHDF1-FMRP interaction in mouse brain tissue. **d**, Illustration of proteins regulating m^6^A-mediated mRNA translation. **d**, Western blotting of mouse hippocampal neurons validating knockout of *Fmr1* gene. Data are mean ± s.e.m. Statistical analysis was performed using unpaired two-tailed *t*-test. Exact *P* values are indicated, and NS denotes *P* values > 0.05.

**Extended Data Fig. 2:**
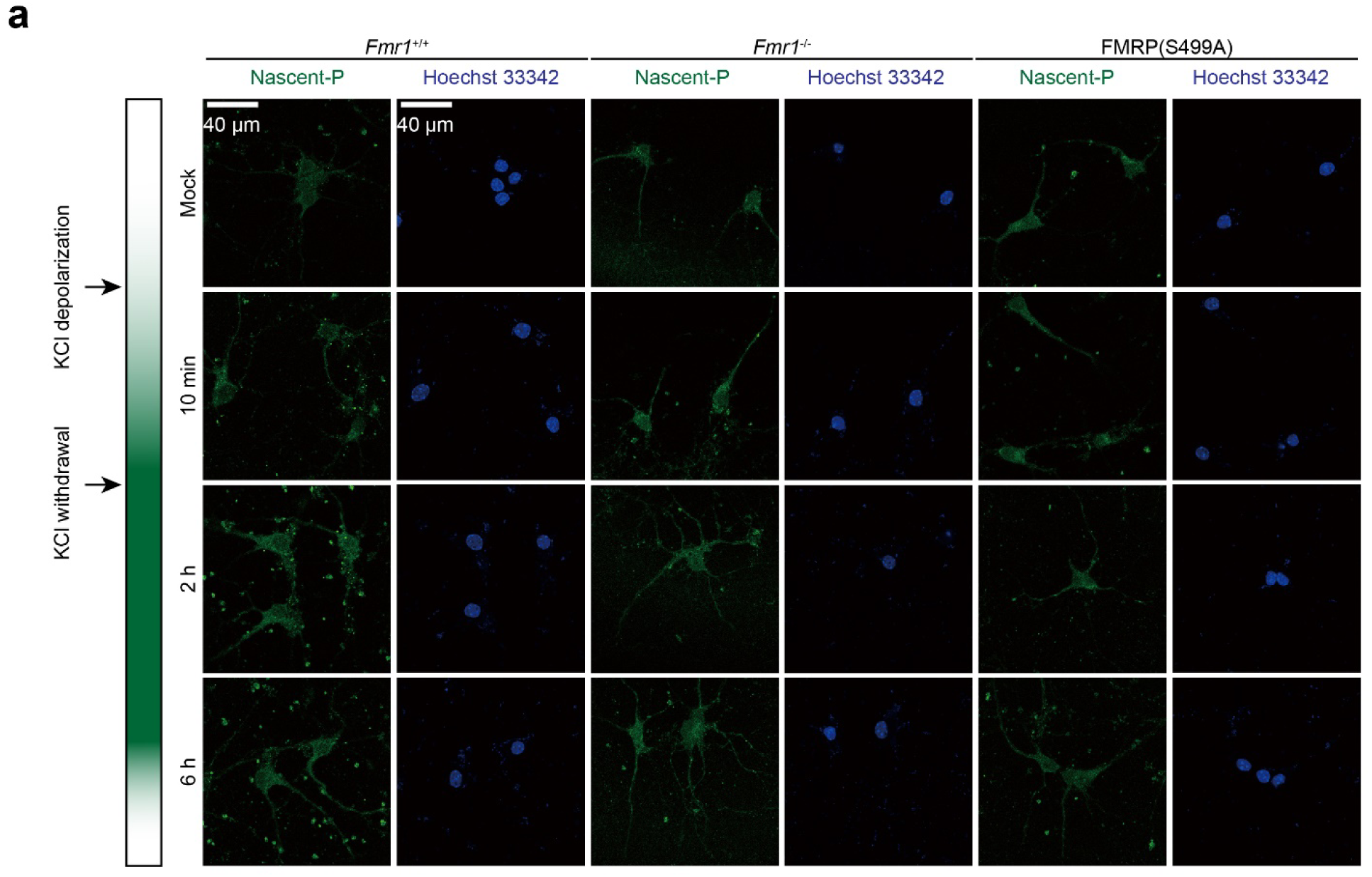
YTHDF1 activation upon neuronal stimulation is dependent on FMRP phosphorylation. **a**, Representative images of protein synthesis rate measured by L-Homopropargylglycine (HPG) incorporation in neurons (n = 6 each condition). Scale bar: 40 µm. Nascent-P: newly synthesized peptides visualized with HPG.

**Extended Data Fig. 3:**
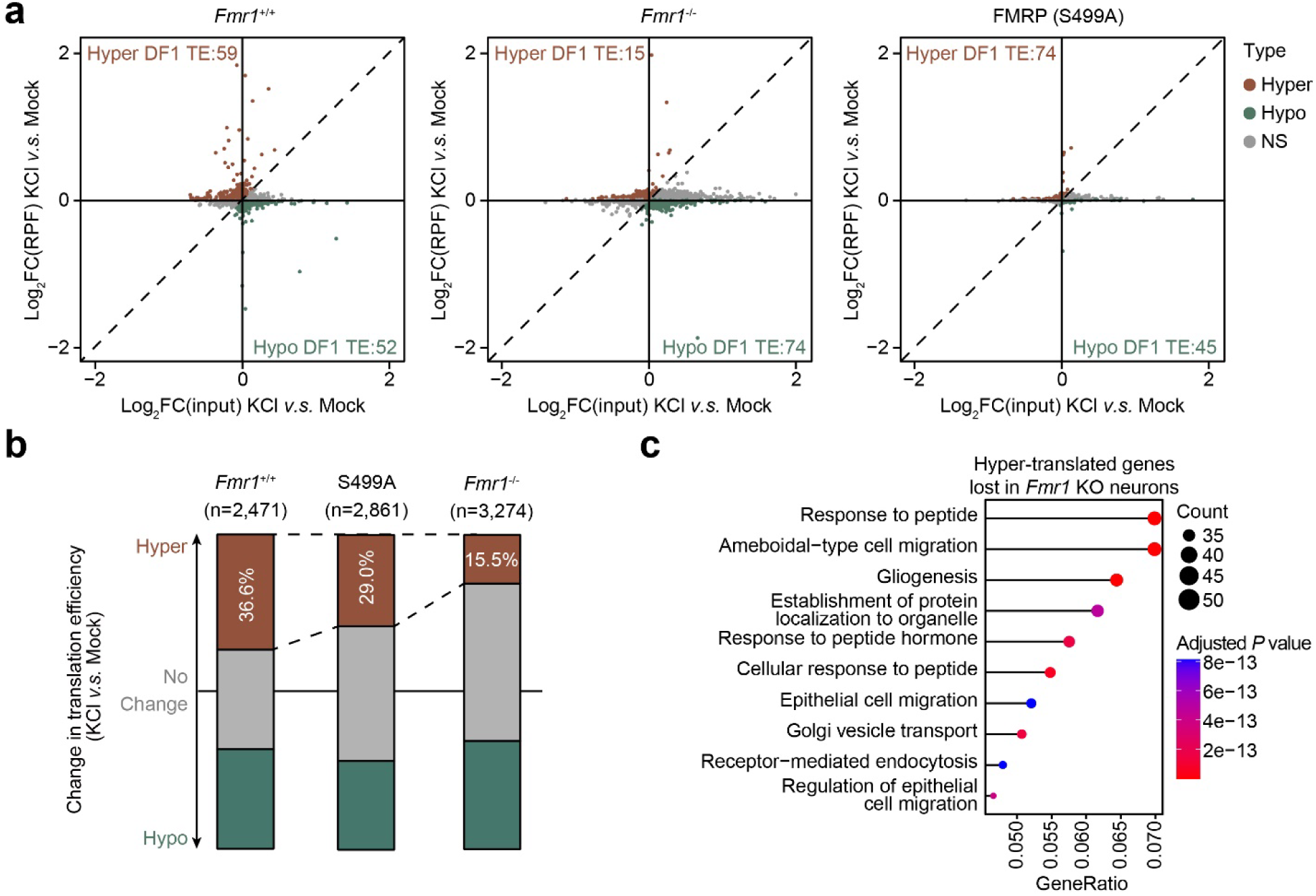
YTHDF1 activation upon neuronal stimulation is dependent on FMRP phosphorylation. **a**, Scatter plots showing changes in RNA translation in cultured neurons upon KCl depolarization. **b**, Bar plots showing the numbers of hyper-translated genes upon KCl depolarization. **c**, Gene ontology enrichments of genes hyper-translated upon KCl depolarization in *Fmr1*^+/+^ neurons but not in *Fmr1*^-/-^ neurons.

**Extended Data Fig. 4:**
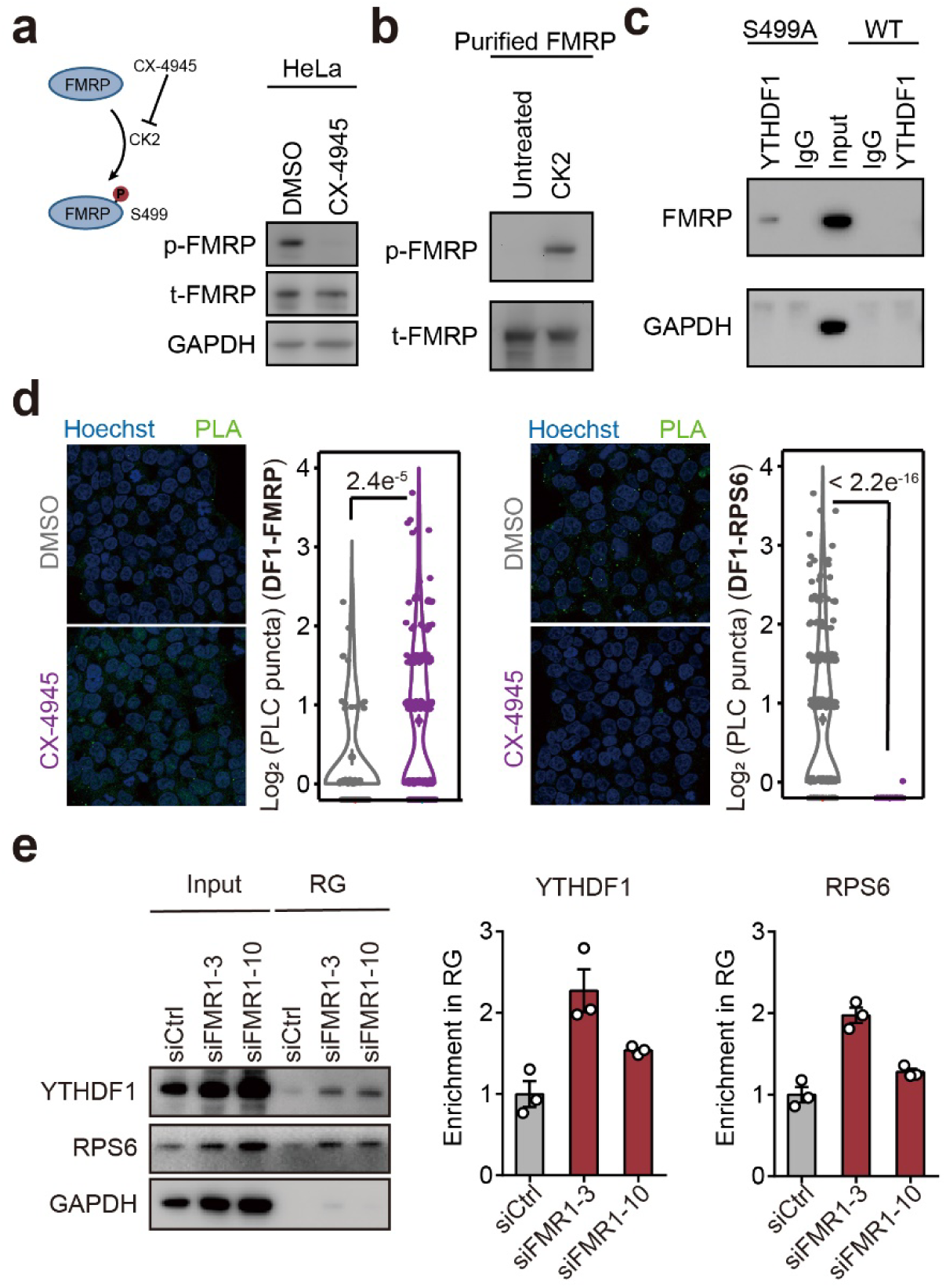
Phosphorylation of FMRP regulates YTHDF1 condensation in HEK293T cells. **a**, Western blot of FMRP S499 phosphorylation in HEK293T cells treated by CX-4945. **b**, Western blot of in vitro phosphorylated recombinant FMRP. **c**, Western blot of association between YTHDF1 and FMRP. FMRP mutant S499A was used to eliminate phosphorylation. **d**, Proximity ligation assay quantifying interactions between YTHDF1/FMRP or YTHDF1/RPS6. **e**, Representative western blot of YTHDF1 and RPS6 in RNA granules (RG) upon FMRP knockdown (n = 3). Western blots are quantified by ImageJ. Relative protein levels were normalized to GAPDH. Statistical analysis was performed with Wilcoxon rank sum test (**d**). Exact *P* values are indicated, and NS denotes *P* values > 0.05.

**Extended Data Fig. 5:**
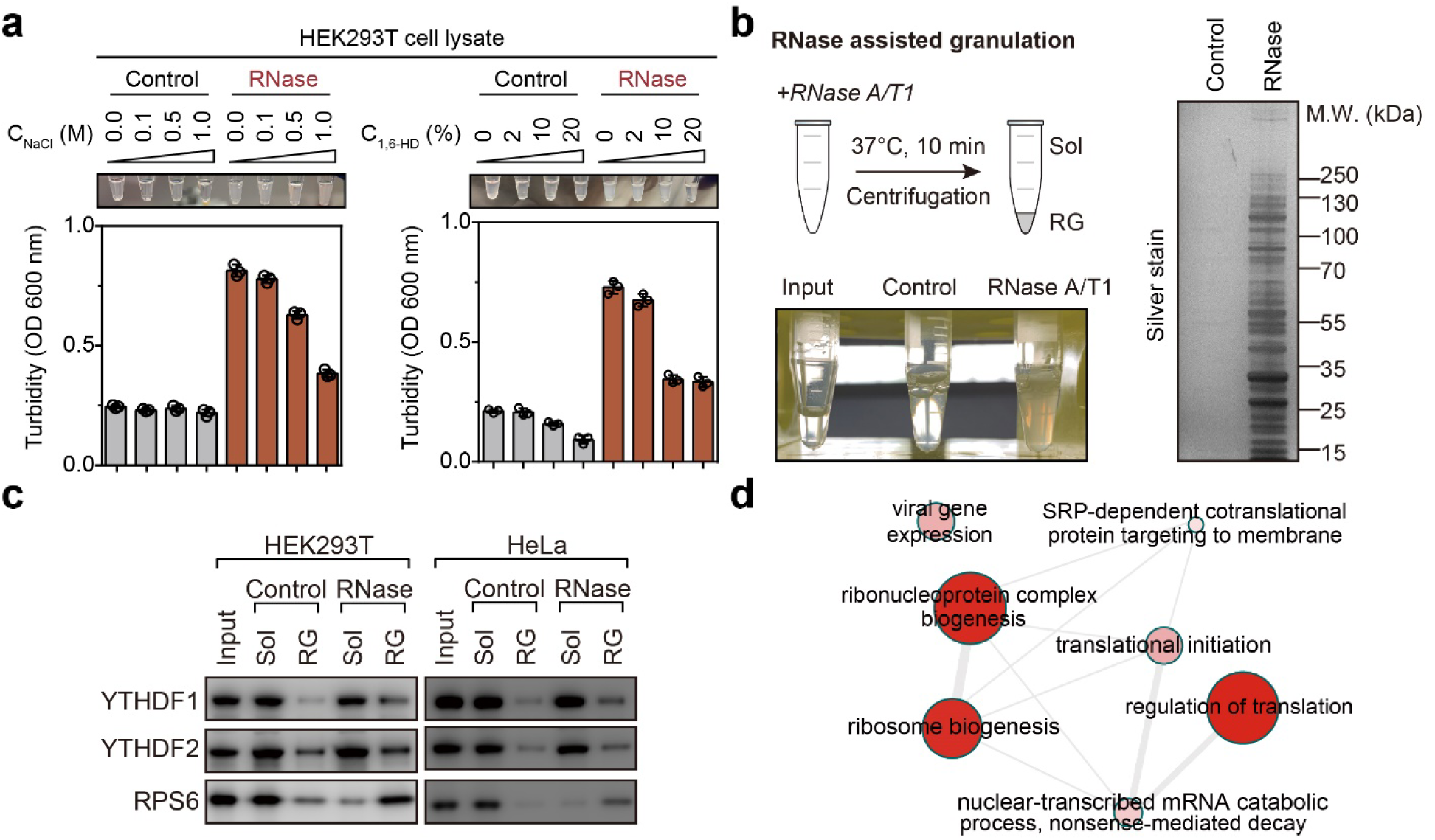
RNase treatment precipitates condensates containing YTHDF1. **a**, Turbidity measurements of HEK293T lysates treated by RNase and high salt (NaCl) or 1,6-hexanediol (1,6-HD). **b**, Analysis of proteins precipitated by RNase treatment. Right: silver stain of proteins present in the condensed phase in response to RNase treatment. **c**, Distribution of YTHDF1, YTHDF2 and RPS6 in the condensed phase in response to RNase treatment. d, Gene ontology terms enriched by condensation-dependent YTHDF1 protein partners.

**Extended Data Fig. 6:**
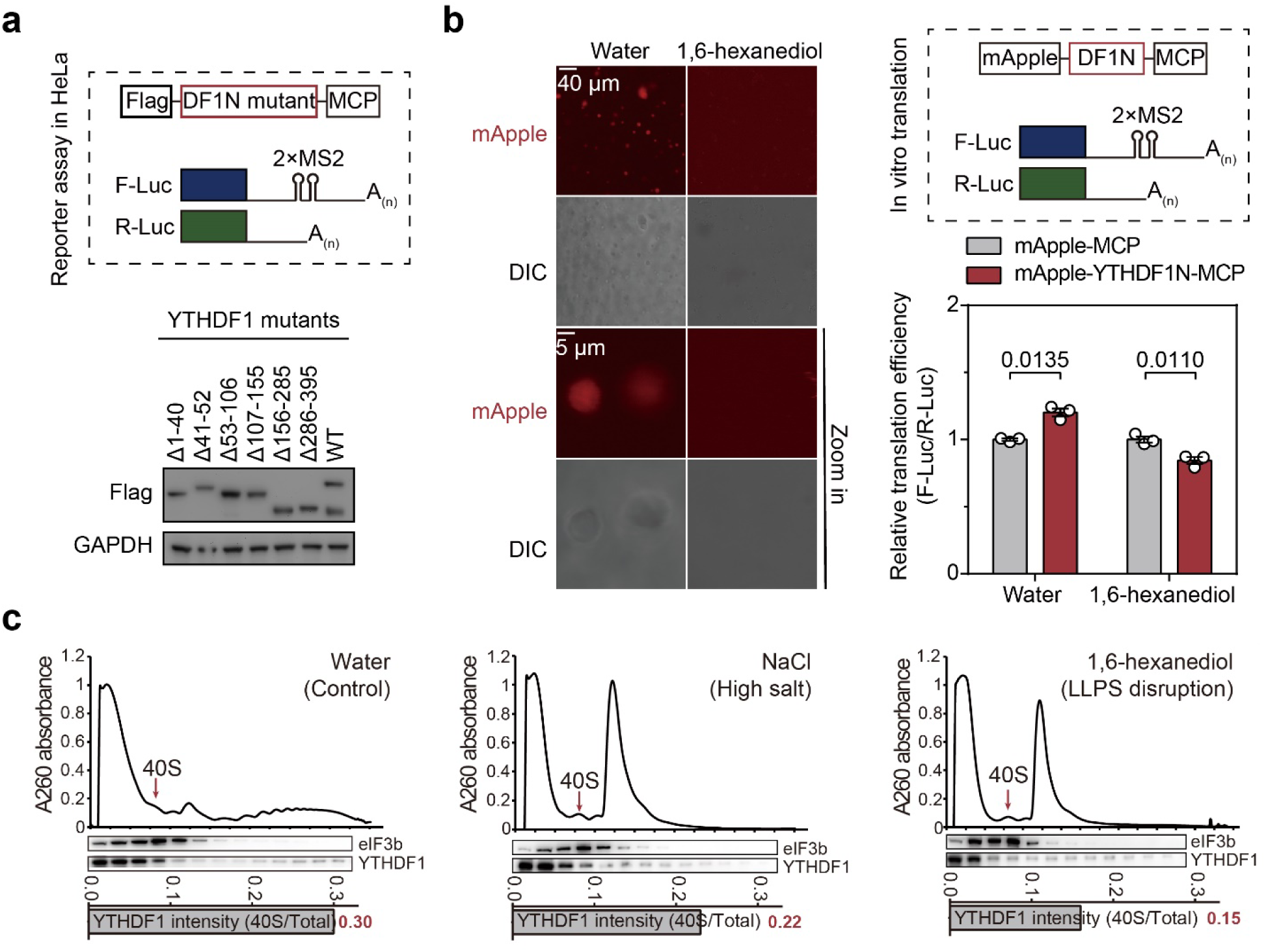
YTHDF1 condensates with ribosome to promote translation. **a**, Illustration of reporter design and western blots validating expression of YTHDF1 mutants. **b**, In vitro translation assay with rabbit reticulocyte lysate (RRL) and YTHDF1 (n = 3 each condition). Scale bar: 40 µm (upper) or 5 µm (lower). Relative translation efficiency of YTHDF1-tethered reporter mRNA was obtained by normalizing firefly luciferase signal to renilla luciferase signal. **c**, Relative distribution of YTHDF1 in polysome fractions. Localization of YTHDF1 in the 40S ribosome fraction is quantified by normalizing intensity of YTHDF1 in 40S ribosome to total intensity. Statistical analysis was performed with unpaired two-tailed *t*-test (**b**). Exact *P* values are indicated, and NS denotes *P* values > 0.05.

**Extended Data Fig. 7:**
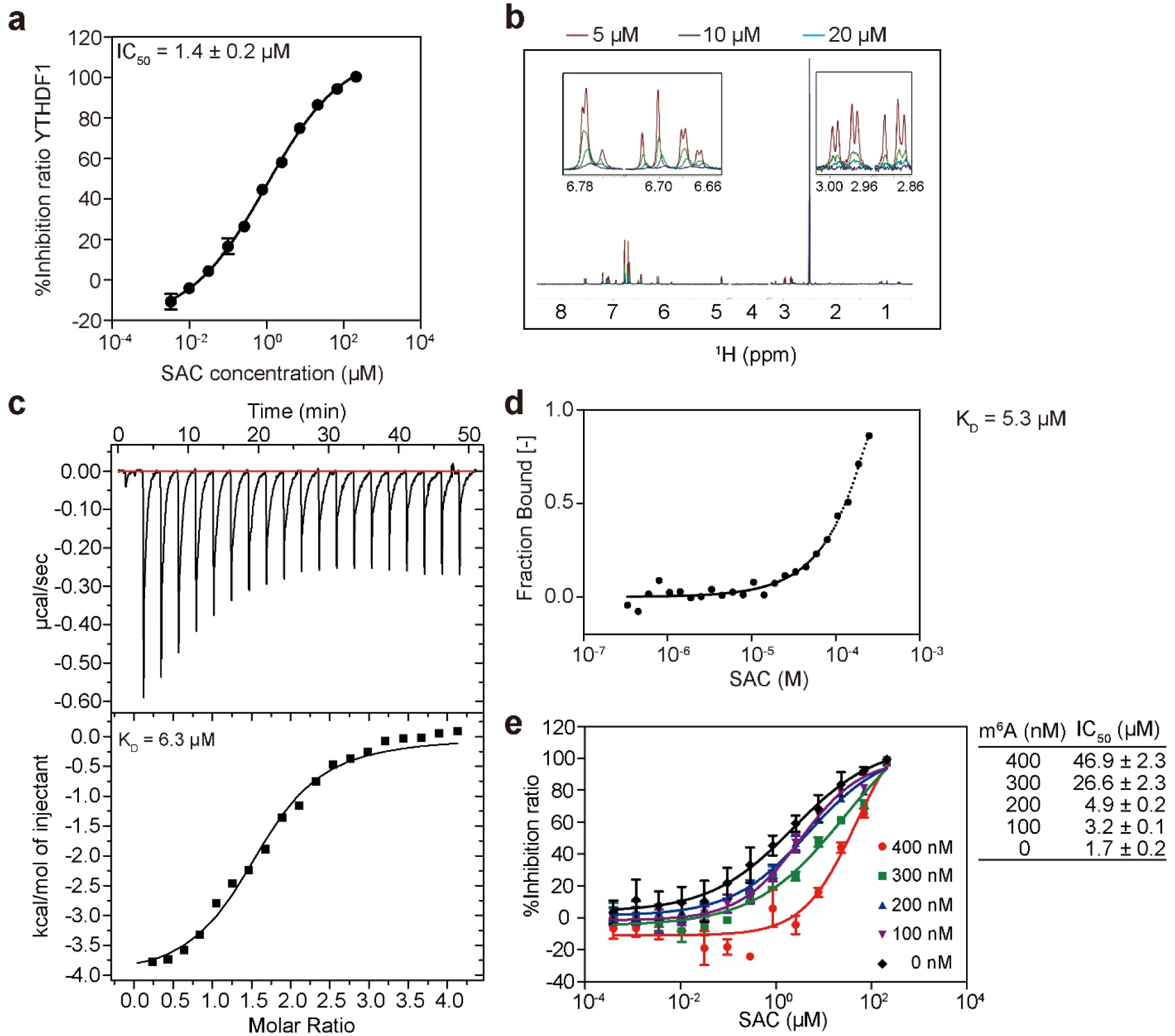
Development of salvianolic acid C (SAC) as a substrate-competitive YTHDF1 inhibitor. a, IC50 value of SAC against YTHDF1 measured by AlphaScreen experiments. b, NMR CPMG binding assay between SAC and YTHDF1. The measurements were conducted under the conditions of 0 µM (red), 5 µM (green), 10 µM (blue) and 20 µM (purple) YTHDF1 protein, respectively. c, ITC analysis between SAC and YTHDF1 at 25 °C. 1 mM SAC was titrated into 50 µM YTHDF1. d, The binding curve of SAC and YTHDF1 obtained via MST assay. e, Competitive binding analysis between SAC and m^6^A-containing RNA oligos. The activities of SAC were summarized in the table, in which m^6^A-containing RNA oligos were shown as m^6^A. (Data are expressed as the mean ± se).

**Extended Data Fig. 8:**
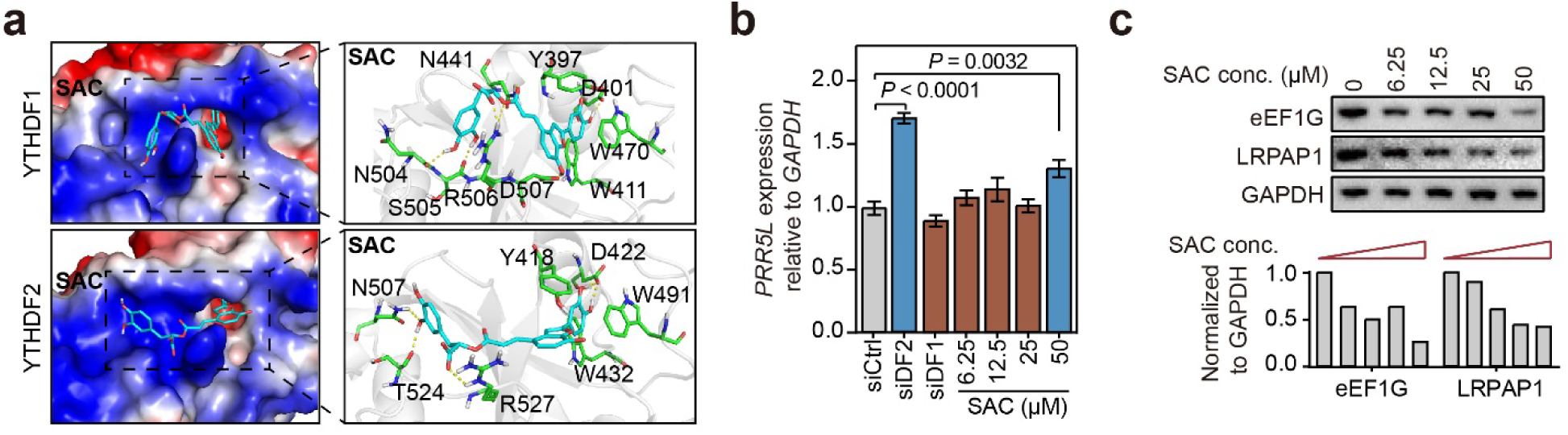
SAC selectively inhibits YTHDF1 at low concentrations. a, Molecular docking of SAC against YTHDF1 or YTHDF2. While SAC occupy the m^6^A-binding pockets of both YTHDF1 and YTHDF2, SAC forms six hydrogen bonds with N411, N504, S505, R506 and D507 of YTHDF1. Especially, SAC forms two hydrogen bonds with R506, which is important to its inhibitory effects. In contrast to YTHDF1, YTHDF2 only forms three hydrogen bonds with SAC (N507, T524 and R527). The more intensive interactions at the entrance of the m^6^A binding pocket of YTHDF1 explain the selectivity. **b**, RNA levels of YTHDF2 target (*PRR5L*) upon gradient SAC treatments by a concentration gradient. *PRR5L* expressions were quantified by normalizing RNA levels to *GAPDH*. **c**, Protein levels of YTHDF1 targets (eEF1G and LRPAP1) upon SAC treatments by a concentration gradient. Protein levels were quantified by western blot intensities and normalized to GAPDH. Data are mean ± se (**a**) or mean ± s.e.m (**b**). Statistical analysis was performed using unpaired two-tailed *t*-test. Exact *P* values are indicated, and NS denotes *P* values > 0.05.

**Extended Data Fig. 9:**
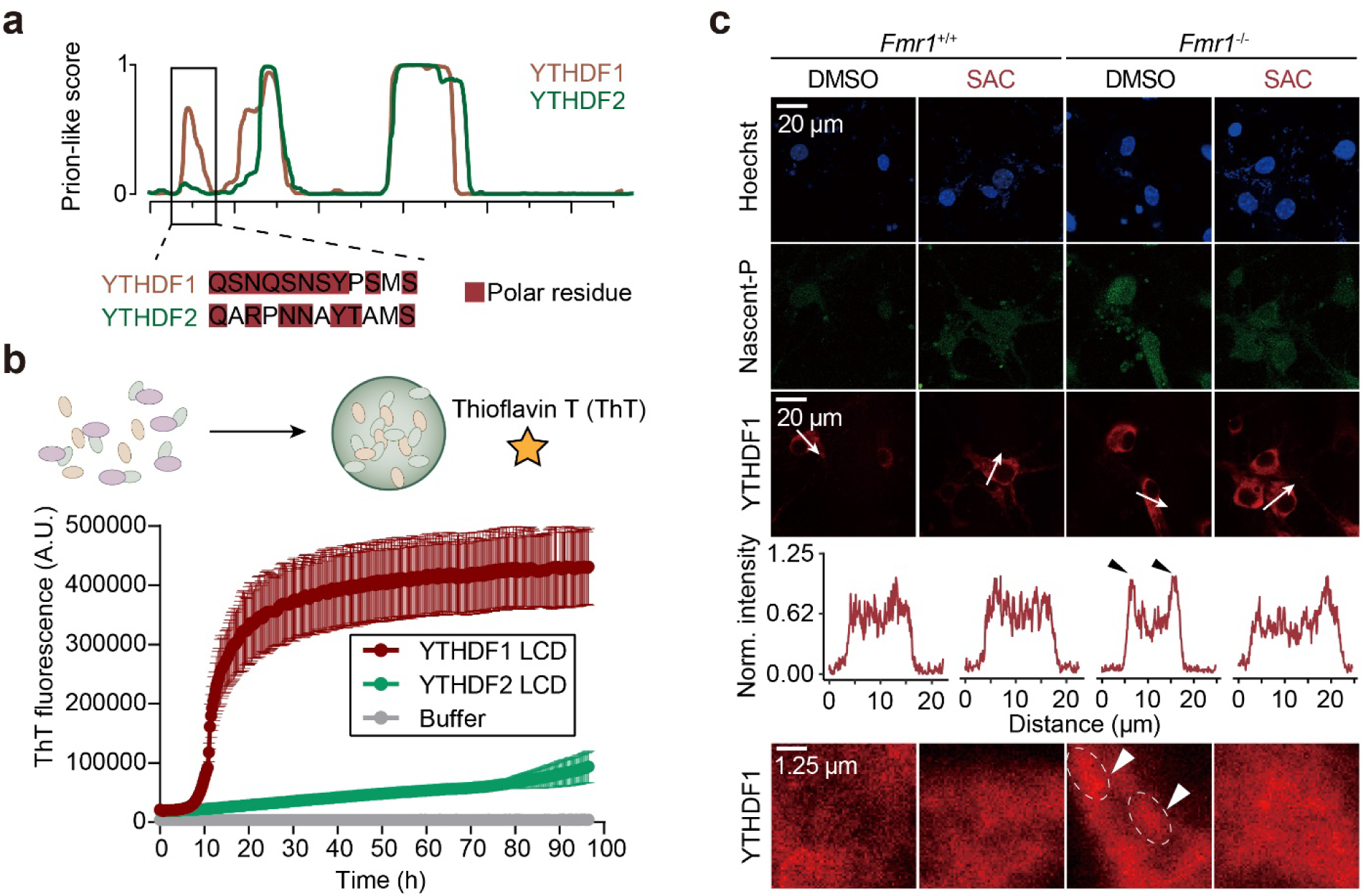
YTHDF1 and YTHDF2 differ in their N-terminal intrinsically disordered domain sequence and exhibit different fiber formation behaviors. **a**, Prion-like score of YTHDF1 and YTHDF2 predicted by PLAAC. **b**, Thioflavin T (ThT) assay of YTHDF1 and YTHDF2 low-complexity domains. Freshly-purified proteins were incubated for 100 hours in native buffer. **c**, Representative images of YTHDF1 cluster and protein synthesis in mouse hippocampal neurons (n = 6 for each condition). Scale bar: 20 µm (upper) or 1.25 µm (lower). Intensity distribution on a line is quantified by ImageJ: Plot profile module.

**Extended Data Fig. 10:**
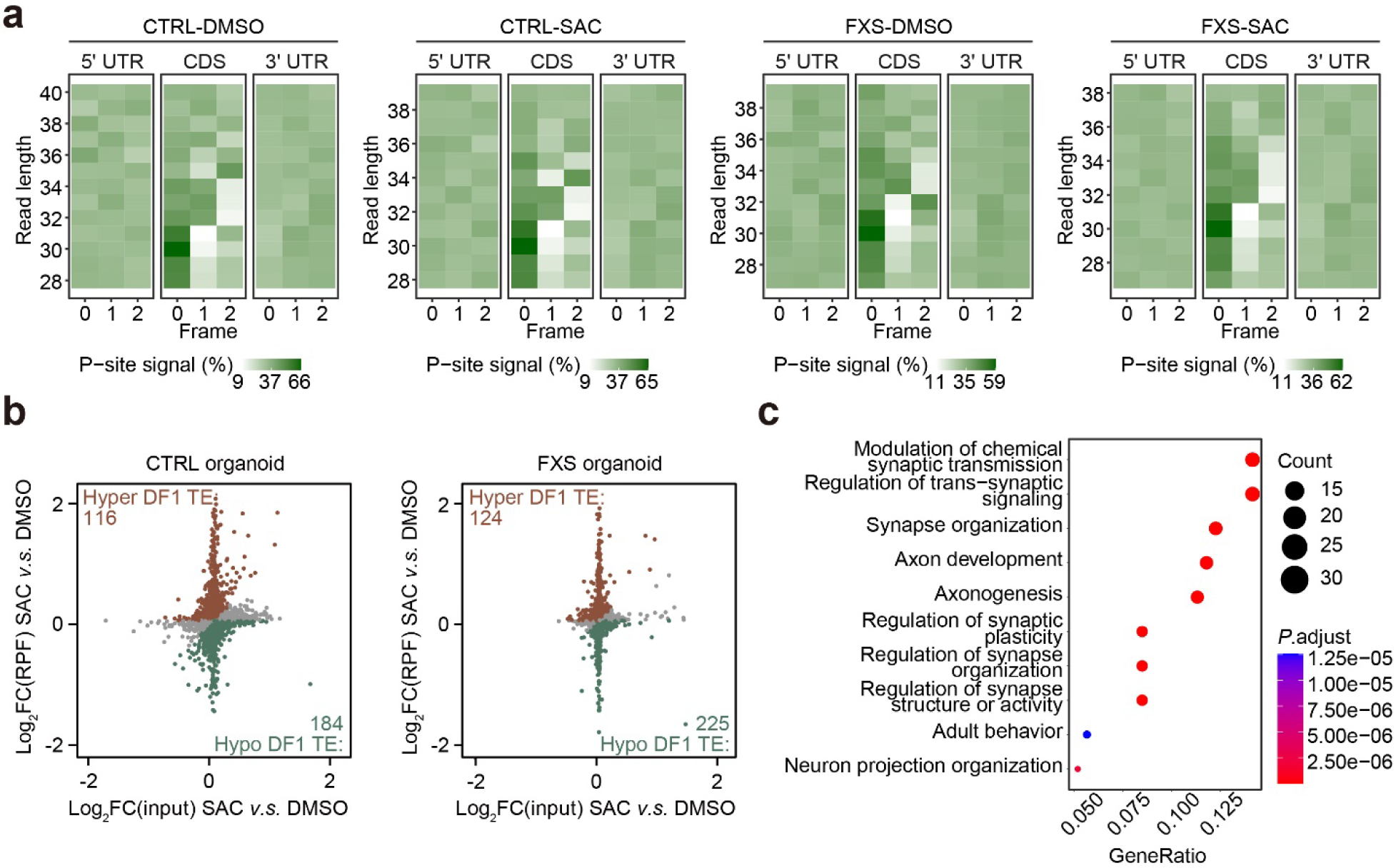
RiboLace measures translation events in organoid models. **a**, Trinucleotide periodicity of P-sites identified with ribosome-protected fragments in individual samples. **b**, Scatter plots showing changes in RNA translation in forebrain organoids upon SAC treatment. **c**, Gene ontology terms enriched by hypo-translated genes in FXS organoids upon SAC treatment.

**Extended Data Fig. 11:**
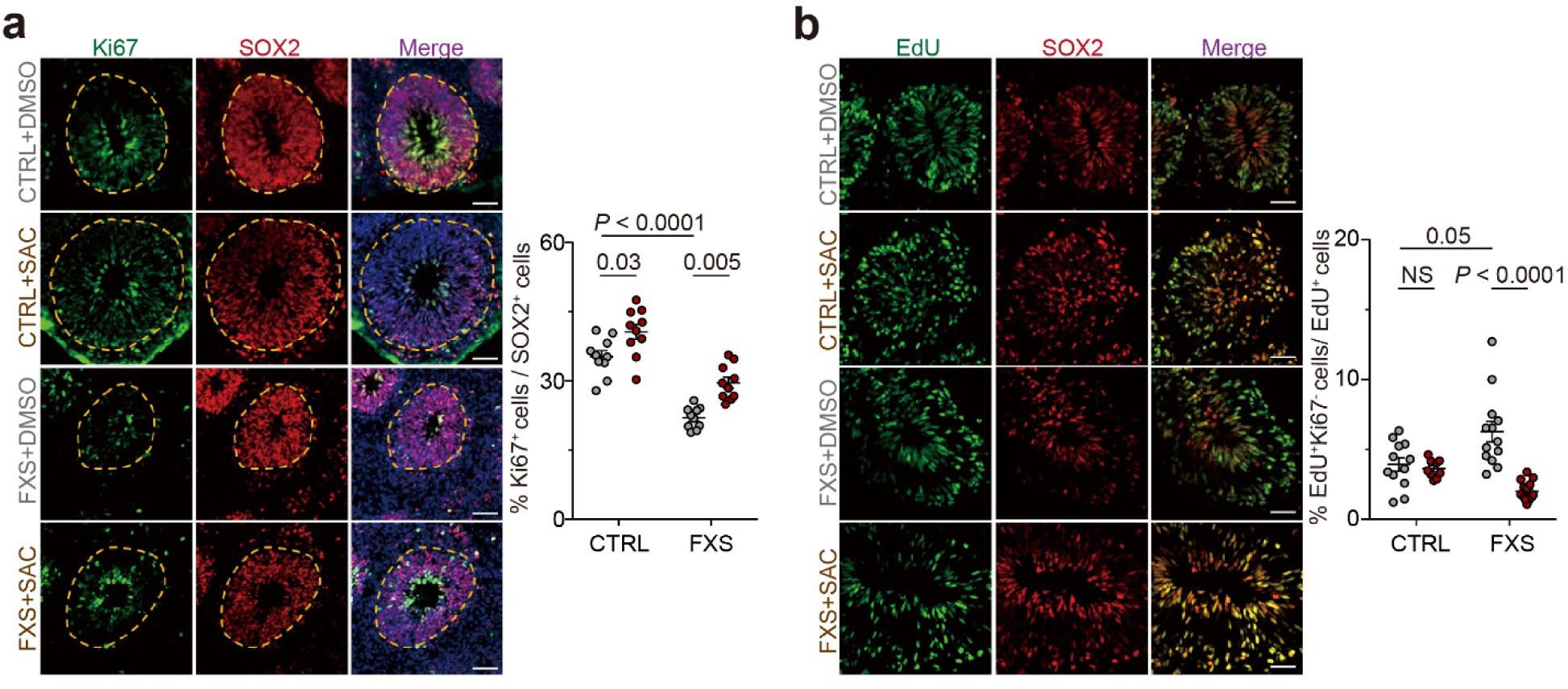
SAC rescues physiopathology in FXS forebrain organoid. **a**, Representative images showing the inhibition of YTHDF1 rescues the reduced NPC proliferation in FXS organoids. **b**, Representative images showing the inhibition of YTHDF1 delays hastened cell cycle exit of cells in FXS organoids. Data are mean ± s.e.m (**b**). Statistical analysis was performed using unpaired two-tailed *t*-test. Exact *P* values are indicated, and NS denotes *P* values > 0.05.

